# Curved adhesions mediate cell attachment to soft matrix fibres in 3D

**DOI:** 10.1101/2023.03.16.532975

**Authors:** Wei Zhang, Chih-Hao Lu, Melissa L. Nakamoto, Ching-Ting Tsai, Anish R. Roy, Christina E. Lee, Yang Yang, Zeinab Jahed, Xiao Li, Bianxiao Cui

## Abstract

Mammalian cells adhere to the extracellular matrix (ECM) and sense mechanical cues through integrin-mediated adhesions^1, 2^. Focal adhesions and related structures are the primary architectures that transmit forces between the ECM and the actin cytoskeleton. Although focal adhesions are abundant when cells are cultured on rigid substrates, they are sparse in soft environments that cannot support high mechanical tensions^3^. Here, we report a new class of integrin-mediated adhesions, curved adhesions, whose formation is regulated by membrane curvature instead of mechanical tension. In soft matrices made of protein fibres, curved adhesions are induced by membrane curvatures imposed by the fibre geometry. Curved adhesions are mediated by integrin ɑVβ5 and are molecularly distinct from focal adhesions and clathrin lattices. The molecular mechanism involves a previously unknown interaction between integrin β5 and a curvature-sensing protein FCHo2. We find that curved adhesions are prevalent in physiologically relevant environments. Disruption of curved adhesions by knocking down integrin β5 or FCHo2 abolishes the migration of multiple cancer cell lines in 3D matrices. These findings provide a mechanism of cell anchorage to natural protein fibres that are too soft to support the formation of focal adhesions. Given their functional importance for 3D cell migration, curved adhesions may serve as a therapeutic target for future development.

## Main

Cells adhere to the extracellular matrix (ECM) through integrin-mediated adhesions to sense mechanical cues and regulate their behaviours in physiology and pathology^1, 2^. Focal adhesions and related structures have been identified as the primary architectures that connect the ECM to the actin cytoskeleton and transmit mechanical forces. Through inside-out signalling, transmembrane integrin receptors are activated and stabilized by force-sensitive intracellular proteins^2^, especially talin^4, 5^. This force-dependent activation is a key regulatory step in the formation of focal adhesions^6^, thus focal adhesions are sensitive to ECM stiffness and cell contractility^7–9^.

Although focal adhesions are numerous and prominent when cells are cultured on 2D rigid substrates, they are much smaller in size^10^, barely discernible^11^, or not observed^12, 13^ in soft 3D matrices. This is partly because talin activation at focal adhesions requires a minimum substrate rigidity of 5kPa^14^, which is higher than the rigidity of most ECM protein fibres^15^. Cell adhesions in 3D are complex and dependent on ECM composition, density, organization, and whether the ECM matrix is synthetic or cell-derived^3^. Despite these complexities, force-dependent integrin activation is still believed to be the key step for cell adhesions in 3D^16^. Interestingly, in conditions where focal adhesions are not observed^12^, adhesion proteins, such as talin, are still important in modulating cell motility in 3D, which raises the possibility that these proteins may participate in alternative adhesion architectures on soft 3D fibres.

Recent studies have identified clathrin-containing integrin adhesions^17^, including flat clathrin lattices^18, 19^, tubular clathrin/AP2 lattices^20^, and reticular adhesions^21^. The clathrin lattices may assist cell adhesions to soft fibres. However, both tubular clathrin/AP2 lattices formed on collagen fibres and flat clathrin lattices formed on soft hydrogels have a lifetime of ∼1 min^18, 20^, which is similar to the lifetime of endocytosis but much shorter than what is expected for stable cell adhesions. More importantly, clathrin-containing adhesions are devoid of mechanotransduction components such as actin and talin^17^. Therefore, clathrin-containing adhesions are likely not the primary architectures responsible for stable cell adhesion and mechanotransduction in 3D.

A key geometrical feature of ECM fibres is their cylindrical and curved surface. It has long been hypothesized that the fibre geometry may play an important role in cell-matrix adhesions by inducing plasma membrane curvature^22^. However, a mechanism that links membrane curvature to integrin adhesion is yet to be identified. In this work, we report the identification of a new architecture of integrin-mediated adhesion - curved adhesion. The formation of curved adhesions is driven by membrane curvature instead of mechanical force and enables cells to adhere to soft fibres in 3D.

### Positive membrane curvature induces selective accumulation of integrin β5

We used a recently developed nanostructure platform to induce well-defined membrane curvatures in live cells^23^. SiO_2_ nanobars were created by etching quartz substrates using electron beam lithography patterning and anisotropic etching. These nanobars are vertically aligned with 200-nm width, 2-µm length, 1.4-µm height, and 5-µm spacing (**Fig. 1a,b** and **Extended Data Fig. 1a**). When adherent cells are cultured on the nanobars, their membranes are deformed to accommodate the geometry of nanobars^24^ (**Fig. 1c**). Membrane wrapping at the two vertical end surfaces and the horizontal top surface of a nanobar results in localized and pre-defined curvatures in the plasma membrane. The flat side walls of nanobars serve as an internal control (**Fig. 1b**). In 2D images that focus on the middle height, due to the 3D-to-2D projection effect, the curvature effect is primarily located at the ends of nanobars with a small contribution from the top surface that distributes uniformly along the length of nanobars. Each quartz substrate has nanopatterned areas (1x1 mm^2^ each) interspersed with control flat areas.

**Fig. 1.**
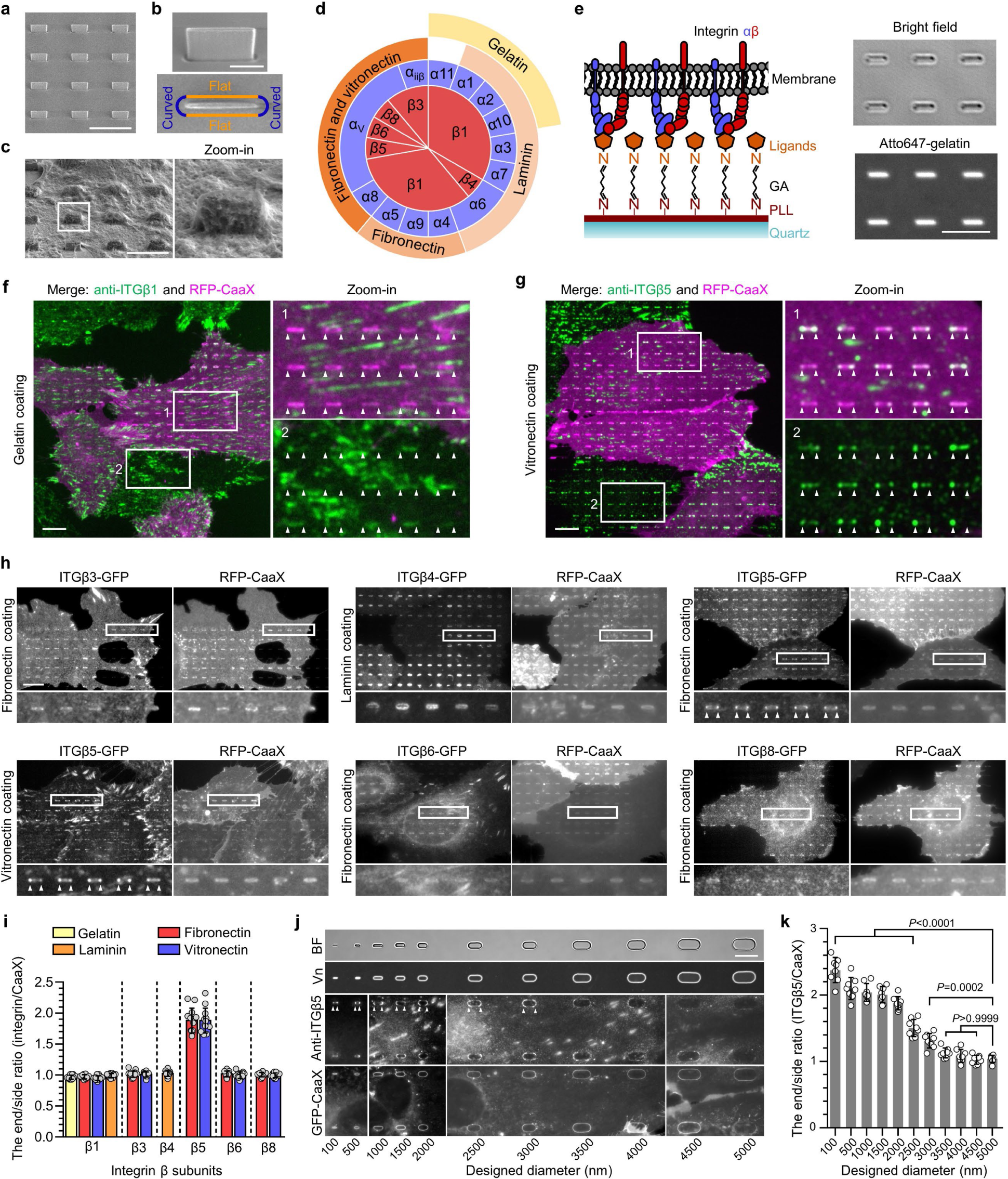
Positive membrane curvature induces selective accumulation of integrin β5. **a,** Scanning electron microscopy (SEM) image of nanobars viewed at 45° angle. **b,** Single nanobar viewed at 45° angle (top) or from the top (bottom). **c,** SEM image showing cell membrane deformation on nanobars. **d,** Chart showing the integrin β subunits probed in this study, with their respective ɑ subunits and ECM ligands. **e,** Left: schematic of ECM ligand-coated surface. Right: bright-field (top) and fluorescence (bottom) images of nanobars coated with Atto647-gelatin. **f,** Anti-ITGβ1 on gelatin-coated nanobar substrate does not show accumulations on nanobars that are visualized by a membrane marker RFP-CaaX transiently expressed in some cells. **g,** Anti-ITGβ5 on vitronectin-coated nanobar substrate shows preferential accumulation at the nanobar ends while RFP-CaaX is relatively evenly distributed on the same nanobars. **h,** Fluorescence images of GFP-tagged β3, β4, β5, β6, and β8 integrins co-expressed with RFP-CaaX on nanobar substrates with their respective ECM protein coatings. Only ITGβ5-GFP shows preferential accumulation at nanobar ends. More images are included in Extended Data Fig. 3a. **i,** Quantifications of integrin β subunits’ curvature preferences by measuring their nanobar end/side ratio and then normalizing the ratios against the end/side ratio of RFP-CaaX at the same locations. Each data point represents a single cell averaging over 30∼160 nanobars. N = 12 cells from two independent experiments. **j,** Probing the curvature range that induces ITGβ5 accumulation using gradient nanobar arrays. First row: bright-field (BF) image of gradient nanobars. Second row: fluorescence image of gradient nanobars coated with Cy3-vitronectin (Vn). Third and fourth rows: anti-ITGβ5 in cells expressing GFP-CaaX on vitronectin-coated gradient nanobars. Accumulations of anti-ITGβ5 at nanobar ends are marked by arrowheads. **k,** Quantification of the end/side ratio of ITGβ5/CaaX on gradient nanobars. N = 8 images for each condition, from two independent experiments. *P* values calculated using one-way analysis of variance (ANOVA) with Bonferroni’s multiple-comparison. Data are mean ± standard deviation (SD). Scale bars: 5 µm (a, c, e); 1 µm (b); 10 µm (f, g, h, j).

In mammals, integrins have 18 ɑ and 8 β subunits, which form 24 distinct ɑβ heterodimers that bind to various ECM ligand proteins^25^. We probed different integrin isoforms by coating nanobar substrates with their respective ECM ligands (**Fig. 1d**). Specifically, we used hydrolyzed collagen (gelatin), laminin, fibronectin, and vitronectin to probe the large β1-containing integrin family. We used fibronectin and vitronectin to probe β3, β5, β6 and β8-containing integrins, and laminin to probe the β4-containing integrin. The leukocyte-specific integrins β2 and β7 that mainly mediate cell-cell adhesions, were not investigated in this study. For surface coating, we first applied poly-L-lysine (PLL) and then crosslinked it to specific ECM proteins using glutaraldehyde (GA). This protocol generates uniform ECM coatings as shown by fluorescent gelatin (**Fig. 1e**). We used immunofluorescence to probe endogenous integrin β1 (ITGβ1) and integrin β5 (ITGβ5), both of which are highly expressed in U2OS cells^26^ (**Extended Data Fig. 2a**), and C-terminal green fluorescent protein (GFP) tagging to probe integrins β3, β4, β6, and β8. We also used GFP tagging to probe integrin β5 and its variants.

On flat areas, endogenous ITGβ1, as well as β3-GFP, β5-GFP, and β6-GFP form focal adhesions on flat substrates coated with their respective ECM ligands, β4-GFP forms hemidesmosomes on laminin, and β8-GFP appears diffusive due to the lack of talin-binding domains (**Extended Data Fig. 2b**). On nanobar areas, anti-ITGβ1 appears in focal adhesions between nanobars but shows little signals on nanobars, which cause membrane deformation visualized by the membrane marker RFP-CaaX (**Fig. 1f**). Interestingly, in addition to being present in focal adhesions between nanobars, anti-ITGβ5 shows selective accumulation at some but not all nanobar ends, which are locations of high membrane curvature, while the co-expressed RFP-CaaX is relatively evenly distributed on the same nanobars (**Fig. 1g**). The distinct distributions of anti-ITGβ1 and anti-ITGβ5 on nanobars are evident in both RFP-CaaX-transfected and non-transfected cells.

On nanobars, β3-GFP, β4-GFP, β6-GFP, and β8-GFP show a relatively even distribution along the nanobar length similar to RFP-CaaX (**Fig. 1h**). ITGβ5-GFP shows selective accumulation at the ends of nanobars on both vitronectin and fibronectin-coated substrates (**Fig. 1h**). More images of anti-ITGβ1 on fibronectin, vitronectin, and laminin, as well β3-GFP, β6-GFP, and β8-GFP on vitronectin-coated nanobars are shown in **Extended Data Fig. 3a** and **3b**. Notably, anti-ITGβ1 does not accumulate at the ends of nanobars coated with any of the ligands. ITGβ5 interacts with the ɑV subunit to form ɑ**_V_**β5 heterodimer for ligand binding. Immunofluorescence with an ɑ**_V_**β5-specific antibody shows that ɑ**_V_**β5, similar to ITGβ5, preferentially accumulates at the nanobar ends (**Extended Data Fig. 3c**), indicating the presence of ɑ**_V_**β5 heterodimers at high curvature locations. In addition to U2OS cells, preferential accumulation of ITGβ5 at the nanobar ends were also observed in many human cell lines including HT1080, A549, U-251, MCF7, HeLa, human mesenchymal stem cells (**Extended Data Fig. 3d**), as well as in mouse MEF cells (**Extended Data Fig. 3e**), indicating that the curvature preference of ITGβ5 is not limited to one specific cell type.

We quantified the curvature preference by measuring the nanobar end/side ratio of integrin β subunits and then normalizing the ratios against the end/side ratio of the membrane marker RFP-CaaX at the same nanobars. The membrane normalization is important to distinguish the curvature effect from the occasional uneven membrane wrapping. The quantifications confirm that ITGβ5 shows a strong preference for high curvature, with an average end/side ratio of ∼1.9 on both vitronectin and fibronectin (**Fig. 1i)**. The end/side ratios of other β subunits are ∼1.0 on their respective ligands, indicating no curvature preference. Since fibronectin is a weak ligand for ɑ**_V_**β5 but a potent ligand for integrin ɑ5β1 that is well expressed in U2OS cells^26^ (**Extended Data Fig. 2a**), we used vitronectin coating for the subsequent ITGβ5 studies unless noted otherwise.

To determine the range of curvature that facilitates the preferential accumulation of ITGβ5, we engineered gradient nanobars whose end-curvature diameters range from 100 nm to 5 µm (**Fig. 1j** and **Extended Data Fig. 1c**). While GFP-CaaX shows that the cell membrane wraps around nanobars of all sizes, anti-ITGβ5 exhibits a clear preference for the ends of thin nanobars, but the curvature preference gradually diminishes as the nanobar width becomes thicker (**Fig. 1j)**. Quantification of the ITGβ5 end/side ratios shows that ITGβ5 prefers sharp curvatures with a curvature diameter of ≤ 3 µm **(Fig. 1k)**.

### Curved adhesions recruit talin-1 and bear low mechanical forces

To determine whether the ITGβ5 accumulations at curved locations participate in cell-matrix adhesions, we used SiO_2_ nanopillar arrays that induce more curvatures per cell than nanobars (**Fig. 2a,b** and **Extended Data Fig. 1b**). ITGβ5-GFP preferentially accumulates at some but not all membrane-wrapped nanopillars as well as in focal adhesions formed between nanopillars on vitronectin-coated substrate, but it appears diffusive on the plasma membrane when the nanopillar substrate is coated with a non-ligand gelatin (**Fig. 2c** and **Extended Data Fig. 4a**). Quantification of ITGβ5-GFP, normalized by RFP-CaaX, shows preferential accumulation of ITGβ5-GFP on vitronectin-coated but not gelatin-coated nanopillars (**Fig. 2d**). We note that, after culturing cells on the substrate for >3 days, ITGβ5-GFP starts to show some accumulations on gelatin-coated nanopillars, likely due to cell-secreted ECM proteins absorbed on the substrate surface. When extracellular Ca^2+^, which is required for the ligand binding of integrins^27^, is sequestered with ethylenediaminetetraacetic acid, ITGβ5-GFP accumulation on vitronectin-coated nanopillars is largely abolished (**Extended Data Fig. 4b,c**). Since ITGβ5 accumulations at curved membranes require ligand binding and participate in cell-ECM adhesions, we term these structures curved adhesions.

**Fig. 2.**
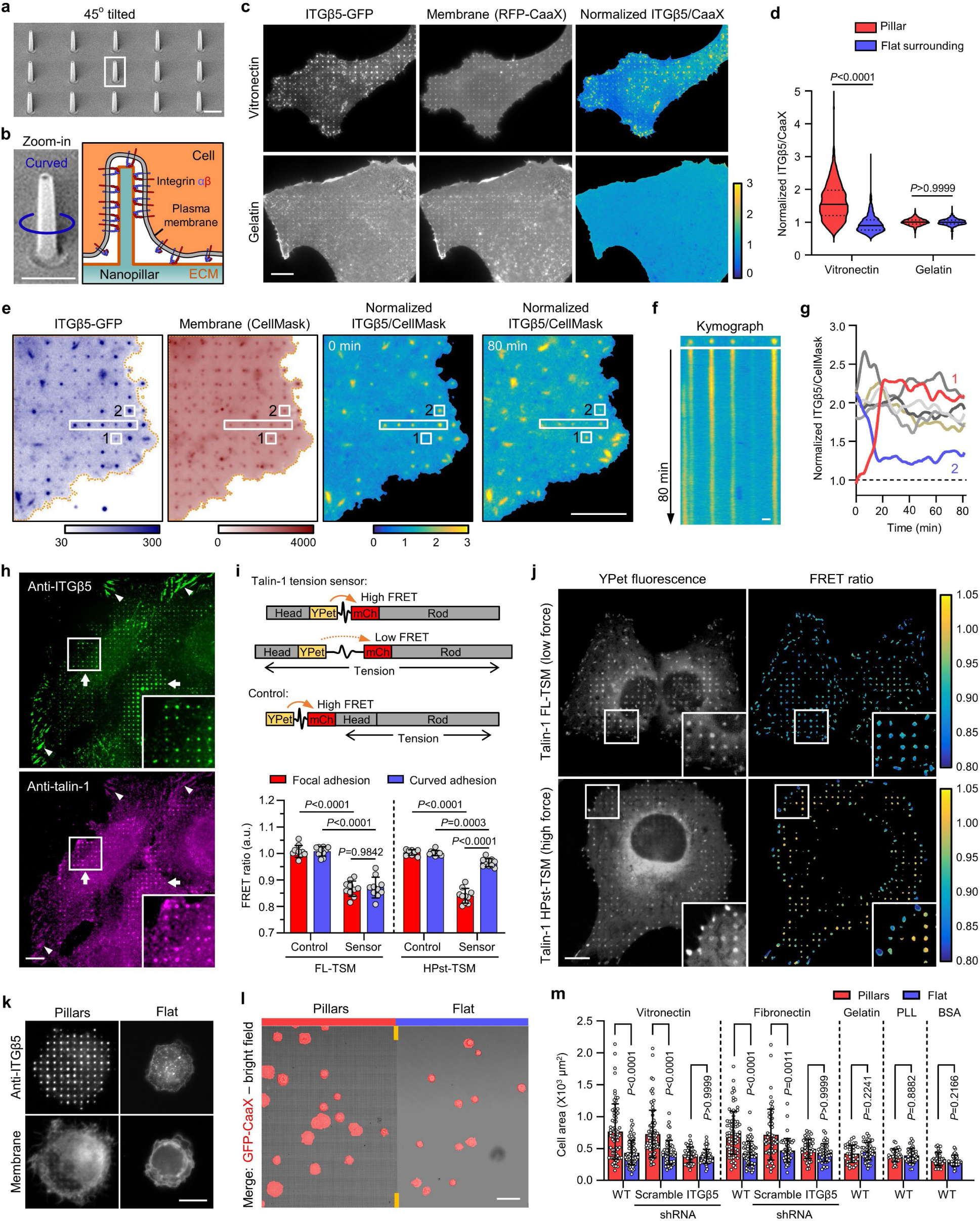
Curved adhesions recruit talin-1 and bear low mechanical forces. **a,** An SEM image of nanopillars. **b,** Left: Zoom in on a single nanopillar in **a**. Right: Schematic of integrin adhesion at a nanopillar. **c,** Fluorescence images showing that vitronectin-coated but not gelatin-coated nanopillars induce ITGβ5-GFP accumulations relative to membrane marker RFP-CaaX. Ratiometric images are shown in the Parula colour scale. **d,** Quantifications of the normalized ITGβ5/CaaX ratio at individual nanopillars and their flat surrounding regions on substrates coated with vitronectin (N= 926 pillars) or gelatin (N = 863 pillars). Medians (lines) and quartiles (dotted lines) are shown. **e,** Live cell imaging of ITGβ5-GFP and RFP-CaaX on vitronectin-coated nanopillar substrate at 15 s/frame for 80 min. Ratiometric images are shown at 0 min and 80 min. **f,** Kymograph of the rectangular box of **e** showing the dynamic of curved adhesions at 4 nanopillars. **g,** Example trajectories of ITGβ5 accumulation at nanopillar locations show that most curved adhesions persist, with a few showing slow assembly (pillar 1) or disassembly (pillar 2). **h,** On vitronectin-coated substrates, endogenous ITGβ5 and talin-1 colocalize in both curved adhesions at nanopillars (arrow) and focal adhesions between nanopillars or on flat areas (arrow heads). **i**, Top: Schematic of talin-1 tension sensors. Bottom: Quantification of the average FRET ratio at focal adhesions and curved adhesions from N = 10 cells for each condition. **j,** Fluorescence (YPet) and ratiometric FRET images (in Parula colour scale) of low-force FL-TSM sensor and high-force HPst-TSM sensor. In the zoom-in window of the HPst-TSM sensor, it is clear that curved adhesions exhibit higher FRET values, and thus lower tensions, than focal adhesions. **k,** ITGβ5 accumulates at vitronectin-coated nanopillars within 30 min after seeding. **l,** Early-stage cell spreading (30 min after plating) on vitronectin-coated nanopillar and flat areas in the same image. **m,** Quantification of the spreading area of N = 30-80 cells in each condition. Data are mean ± SD (**i**, **m**). *P* values calculated using Kruskal-Wallis test with Dunn’s multiple-comparison (**d, m-**vitronectin, fibronectin coatings), one-way analysis of variance (ANOVA) with Tukey’s multiple-comparison (**i**), Mann-Whitney test (**m-**Gelatin coating) or t-test (**m-**PLL, BSA coatings). Scale bars: 1 µm (**a, b, f**); 10 µm (**c, e, h, j, k**); 50 µm (**l**).

Dynamic measurements of ITGβ5-GFP and RFP-CaaX show that curved adhesions are stable adhesions. The RFP-CaaX channel shows that the cell membrane similarly wraps around all nanopillars, while ITGβ5-GFP selectively accumulates on ∼30% of these nanopillars (**Fig. 2e**). As shown in **Fig. 2f** and **Supplementary Video 1**, the vast majority of ITGβ5-GFP-marked curved adhesions (24 out of 28 nanopillars in the image) persist for >80 min of imaging time. Nevertheless, assembly (2 nanopillars, example red trace) and disassembly (3 nanopillars, example blue trace) of curved adhesions occur at a few nanopillars without significant changes in their membrane wrapping (**Fig. 2g** and **Extended Data Fig. 4d**). We note that both the assembly and the disassembly processes are slow and gradual, occurring on the time scale of >10 min.

To explore whether curved adhesions can bear mechanical forces, we first examined whether they recruit talin-1, a key component that mediates force transmission between integrins and the actin cytoskeleton^5^. Co-immunostaining of ITGβ5 and talin-1 shows that talin-1 colocalizes with ITGβ5 on vitronectin-coated nanopillars and selectively accumulates at those nanopillars where ITGβ5 assembles (**Fig. 2h**). In the same image, talin-1 also colocalizes with ITGβ5 in focal adhesions as expected. On the other hand, co-immunostaining of ITGβ1 and talin-1 shows that ITGβ1 does not accumulate at vitronectin-coated nanopillars where talin-1 accumulates (**Extended Data Fig. 4e**). These data demonstrate that that ITGβ5, but not ITGβ1, recruits talin-1 to curved adhesions formed on vitronectin-coated nanopillars. Furthermore, on gelatin-coated substrates, ITGβ1 and talin-1 strongly colocalize in focal adhesions on flat areas, but neither ITGβ1 nor talin-1 accumulates on gelatin-coated nanopillars (**Extended Data Fig. 4f**). With gelatin coating, ITGβ5 appears diffusive and shows no accumulation at nanopillar locations (**Extended Data Fig. 4g**), confirming that curved adhesions do not form on gelatin-coated substrates.

To further investigate mechanical forces in curved adhesions, we employed two genetically encoded and Förster resonance energy transfer (FRET)-based talin-1 tension biosensors, where a tension sensor module (TSM) is inserted between the head domains and the rod domains of talin-1 ^28, 29^ (**Fig. 2i**). When talin-1 is under mechanical forces, the TSM can be stretched, which result in a reduced FRET efficiency. As a negative control, the TSM module is linked to the N-terminus of talin-1, which is not affected by the forces exerted on talin-1. Specifically, FL-TSM is sensitive to low forces with a maximal sensitivity 3-5 pN, while HPst-TSM is sensitive to high forces with a maximal sensitivity 9-11 pN ^29^. Both FL-TSM and HPst-TSM accumulate in curved adhesions at nanopillars as well as in focal adhesions (YPet fluorescence in **Fig. 2j**), a pattern similar to endogenous talin-1.

To compare the mechanical forces involved in curved adhesions and focal adhesions, we quantified the FRET ratio using ratiometric imaging as previously described^28, 29^. The low force FL-TSM sensor shows similar FRET ratios in curved adhesions and focal adhesions, which are significantly lower than the FRET ratios exhibited by its control construct (**Fig. 2i,j**), indicating that the FL-TSM sensor is fully stretched in both adhesion architectures. Interestingly, when the high force HPst-TSM sensor is used, focal adhesions show significantly lower FRET ratios than curved adhesions, indicating that the HPst-TSM sensor is fully stretched in focal adhesions, but only partially stretched in curved adhesions. Together, these results indicate that curved adhesions bear mechanical forces, but the tension along talin-1 molecules in curved adhesions is lower than the tension along talin-1 molecules in focal adhesions.

Besides bearing forces, curved adhesions also promote early-stage cell spreading, a characteristic of functional cell adhesions^30^. Immunofluorescence shows strong ITGβ5 accumulation at vitronectin-coated nanopillars 30 minutes after cell plating, which is before the formation of focal adhesions (**Fig. 2k** and **Extended Data Fig. 5a**). On the other hand, cells largely appear small and round on flat areas. Additionally, the plasma membrane marker GFP-CaaX shows that cell areas on nanopillar areas are visibly larger than those on flat surfaces in the same image (**Fig. 2l**). Similar results were also observed for cells cultured on fibronectin-coated substrates, but not on gelatin, PLL, or BSA-coated substrates (quantifications in **Fig. 2m** and example images in **Extended Data Fig. 5b**). ITGβ5-knockdown (KD) cells show no nanopillar-assisted early-stage cell spreading on either vitronectin or fibronectin-coated substrates (**Extended Data Fig. 5a,c**; KD efficiency verified in Fig. 6). Therefore, nanopillar-assisted early-stage cell spreading is specific to ITGβ5 and its ECM ligands.

### Curved adhesions involve a subset of adhesion proteins and require the juxtamembrane region of ITGβ5’s cytoplasmic domain

To explore whether curved adhesions contain other adhesion proteins besides talin-1, we examined the spatial correlations between ITGβ5 and paxillin, vinculin, phospho-focal adhesion kinase (Tyr397, pFAK), and zyxin. On flat areas, these focal adhesion proteins show strong colocalization with ITGβ5-marked focal adhesion patches (**Extended Data Fig. 6a-d**). However, vinculin and pFAK are absent from curved adhesions marked by ITGβ5 accumulations at nanopillars (**Fig. 3a** and **Extended Data Fig. 6e**), even though they colocalize with ITGβ5 in focal adhesions formed between nanopillars in the same images. On the other hand, paxillin and zyxin show clear colocalizations with ITGβ5, similar to talin-1, in both curved adhesions and focal adhesions (**Extended Data Fig. 6f,g**). Quantifications of Pearson’s correlation coefficients at vitronectin-coated nanopillars suggest that curved adhesions contain a subset of adhesion proteins including talin-1, paxillin, and zyxin, but not vinculin or pFAK (**Fig. 3b**). Additionally, curved adhesions are not linked to large bundles of stress fibres, but are connected to a population of stable F-actin assembled at nanopillars (data and discussions in **Extended Data Fig. 7**).

**Fig. 3.**
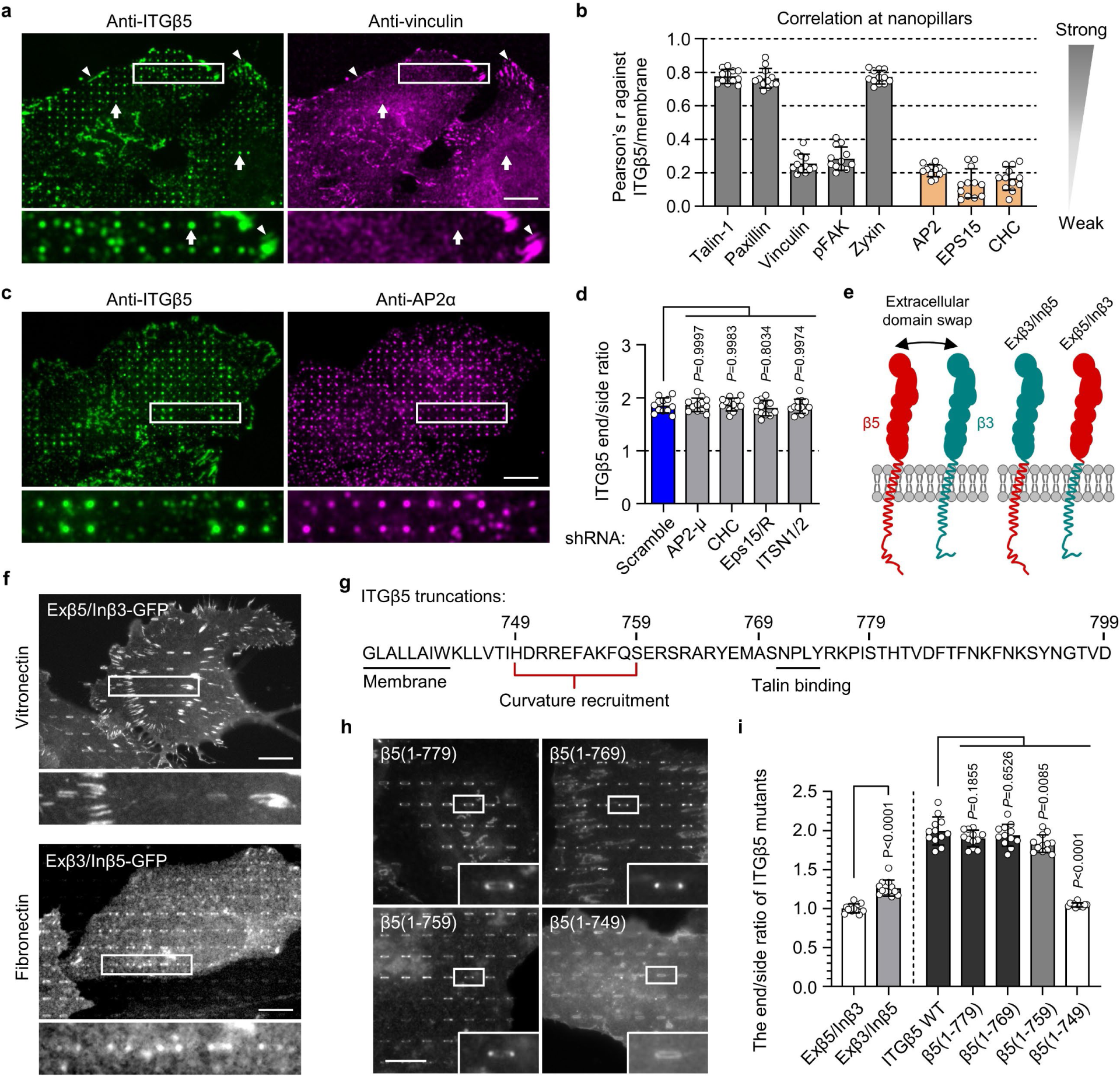
Curved adhesions involve a subset of adhesion proteins and require the juxtamembrane region of ITGβ5 cytoplasmic domain. **a**, Vinculin colocalizes with ITGβ5 in focal adhesions (arrow heads) on flat areas but is absent from curved adhesions (arrows) at nanopillars. **b**, Pearson correlation coefficients between ITGβ5 and focal adhesion proteins (grey bars) or between ITGβ5 and clathrin-mediated endocytic proteins (orange bars) at nanopillars. N = 12 cells for each condition with each cell represents the average of over 100 nanopillars. **c,** Both ITGβ5 and AP2-ɑ preferentially localize at nanopillars but their intensities are not correlated. Zoom-in images show that nanopillars with high AP2 intensities often have lower ITGβ5 intensities. **d,** Quantifications of ITGβ5 accumulation in curved adhesions (end/side ratio on nanobars) upon shRNA knockdown of different endocytic proteins. **e**, Illustration of the construction of chimeric Exβ5/Inβ3 and Exβ3/Inβ5 proteins. **f,** Exβ5/Inβ3 forms prominent focal adhesions but does not accumulate at the ends of nanobars, while Exβ3/Inβ5 shows curvature preference for nanobar ends. **g,** Sequence of the ITGβ5 cytoplasmic domain and the truncation sites. **h,** Fluorescence images of GFP-tagged ITGβ5 truncations (1-779), (1-769), (1-759), and (1-749) on vitronectin-coated nanobars. All truncations, except β5(1-749), show curvature preference for nanobar ends. **i,** Quantification of the curvature preferences of chimeric, wild type (WT), and truncated ITGβ5 by measuring their nanobar end/side ratio. N = 12 cells for each condition. Data are mean ± SD, from two independent experiments. *P* values calculated using one-way ANOVA with Bonferroni’s multiple-comparison (**d**, **i-**WT vs. truncations) or t-test (**i-** Exβ5/Inβ3 vs. Exβ3/Inβ5). Scale bars: 10 µm.

Next, we explored whether curved adhesions are related to clathrin-containing adhesions such as flat clathrin lattices (FCL) and reticular adhesions^17^. Clathrin-containing structures have been reported to recruit β5 as well as β1 and β3 integrins^17–20, 31^. Clathrin and its adaptor AP2 have also been shown to accumulate at nanopillar-induced membrane curvatures for enhanced endocytosis on gelatin-coated nanopillars^24^. Here, immunofluorescence shows that both ITGβ5 and AP2 accumulate at some vitronectin-coated nanopillars (**Fig. 3c**). However, a closer examination shows that their accumulations are not spatially correlated, e.g., nanopillars with high β5 intensities often have low AP2 intensities and vice versa (**Fig. 3c** and quantification in **3b**). Even when both ITGβ5 and AP2 accumulate at the same nanopillars, expansion microscopy imaging of the cell-nanopillar interface^32^ shows that they are not spatially overlapping (**Extended Data Fig. 8a**). Similarly, ITGβ5-GFP accumulations at nanopillars do not correlate with the anti-clathrin heavy chain (anti-CHC) or EPS15-RFP, another clathrin adaptor protein implicated in clathrin-containing adhesions (**Extended Data Fig. 8b,c** and quantifications in **Fig. 3b**). Furthermore, the shRNA knockdown of clathrin heavy chain CHC), AP2 μ subunit (AP2-μ), adaptor proteins EPS15/R, or intersectin1/2 does not affect the curvature preference of ITGβ5-GFP (**Extended Data Fig. 9a,b** and quantifications in **Fig. 3d**). These proteins are key proteins involved in clathrin-containing adhesions. Therefore, curved adhesions are not clathrin-containing adhesions. This is further supported by the presence of talin-1, paxillin, zyxin and F-actin in curved adhesions (**Fig. 2h**; **Extended Data Fig. 6f,g** and 7), because clathrin-containing adhesions are devoid of talin-1, paxillin, F-actin or other mechanotransduction components^17^.

To understand the molecular mechanisms, we probed why ITGβ5, but not its highly homologous isoform ITGβ3, participates in curved adhesions. Two chimeric proteins Exβ5/Inβ3 and Exβ3/Inβ5 were engineered by swapping the extracellular domains of ITGβ3 and ITGβ5 (**Fig. 3e**). Exβ5/Inβ3-GFP forms strong focal adhesions on vitronectin-coated substrates but does not accumulate in curved adhesions at the ends of nanobars (**Fig. 3f** and quantification in **Fig. 3i**). Conversely, though Exβ3/Inβ5-GFP is not well expressed, it shows clear curvature preference on fibronectin-coated nanobars (**Fig. 3f** and quantification in **Fig. 3i**), indicating that the transmembrane domain and the cytoplasmic domain of ITGβ5, but not its extracellular domains, are crucial for the formation of curved adhesions.

We then sequentially truncated the ITGβ5 cytoplasmic domain from the C-terminus, resulting in β5(1-779), β5(1-769), β5(1-759) and β5(1-749) (**Fig. 3g**). We find that, except for β5(1-749), all truncated ITGβ5 variants show curvature sensitivity with preferential accumulations at the ends of nanobars (**Fig. 3h** and quantification in **Fig. 3i**). With the cytoplasmic domain mostly deleted, β5(1-749) behaves like a membrane marker and does not display any curvature preference. It is worth noting that β5(1-769) and β5(1-759), where the NPLY talin-binding motif^4^ is deleted, do not localize to focal adhesions. Nevertheless, at low expression levels, both β5(1-769) and β5(1-759) show clear curvature preference (**Fig. 3h**). Therefore, the curvature preference of ITGβ5 is not dependent on talin binding but requires its intracellular juxtamembrane region.

### Curved adhesions require a curvature-sensing protein FCHo2

When we were probing various proteins related to clathrin-mediated endocytosis, we found that shRNA knockdown of FCHo1/2, two homologous early-stage endocytic proteins, significantly reduced ITGβ5 accumulations at nanobar ends (**Extended Data Fig. 9c**). FCHo1/2 are intrinsic curvature sensing proteins harbouring an N-terminal F-BAR domain and are essential for the initiation of clathrin-mediated endocytosis^33^. FCHo1/2 have not been reported to be involved in cell adhesions. Indeed, FCHo1/2 are not identified in adhesome of either focal adhesions^34^ or clathrin-containing reticular adhesions^21^.

RFP-FCHo2 shows strong and selective colocalization with ITGβ5-GFP at nanopillars (arrows) but does not colocalize with ITGβ5 in focal adhesions (arrow heads) or clathrin-containing adhesions (thin arrows) in the same cell (**Fig 4a**). At nanopillars, the intensity of FCHo2 is positively correlated with the intensity of ITGβ5 with a Pearson’s correlation coefficient of ∼0.6 (**Fig 4b**). Live cell imaging shows that FCHo2 accumulations are strong and stable in curved adhesions marked by ITGβ5 (**Fig 4c** and **Supplementary Video 2**). Some nanopillars without ITGβ5 also show weak FCHo2 accumulations exhibiting frequent and dynamic assembly and disassembly (**Fig. 4c,d**), which are likely involved in endocytosis or other dynamic processes. Anti-ITGβ5 staining shows significantly reduced curved adhesions in U2OS cells transfected with FCHo2 shRNAs, marked by BFP expression, as compared with non-transfected cells in the same image (**Fig 4e**). Quantification of many cells confirms that FCHo2 knockdown significantly reduces ITGβ5 accumulations at nanopillars (**Fig 4g**). Moreover, when ITGβ5 is knocked down with shRNA, FCHo2 accumulation at nanobar ends is also significantly reduced (**Extended Data Fig. 10a,b**). These data suggest that FCHo2 is a stable and essential component of curved adhesion.

**Fig. 4.**
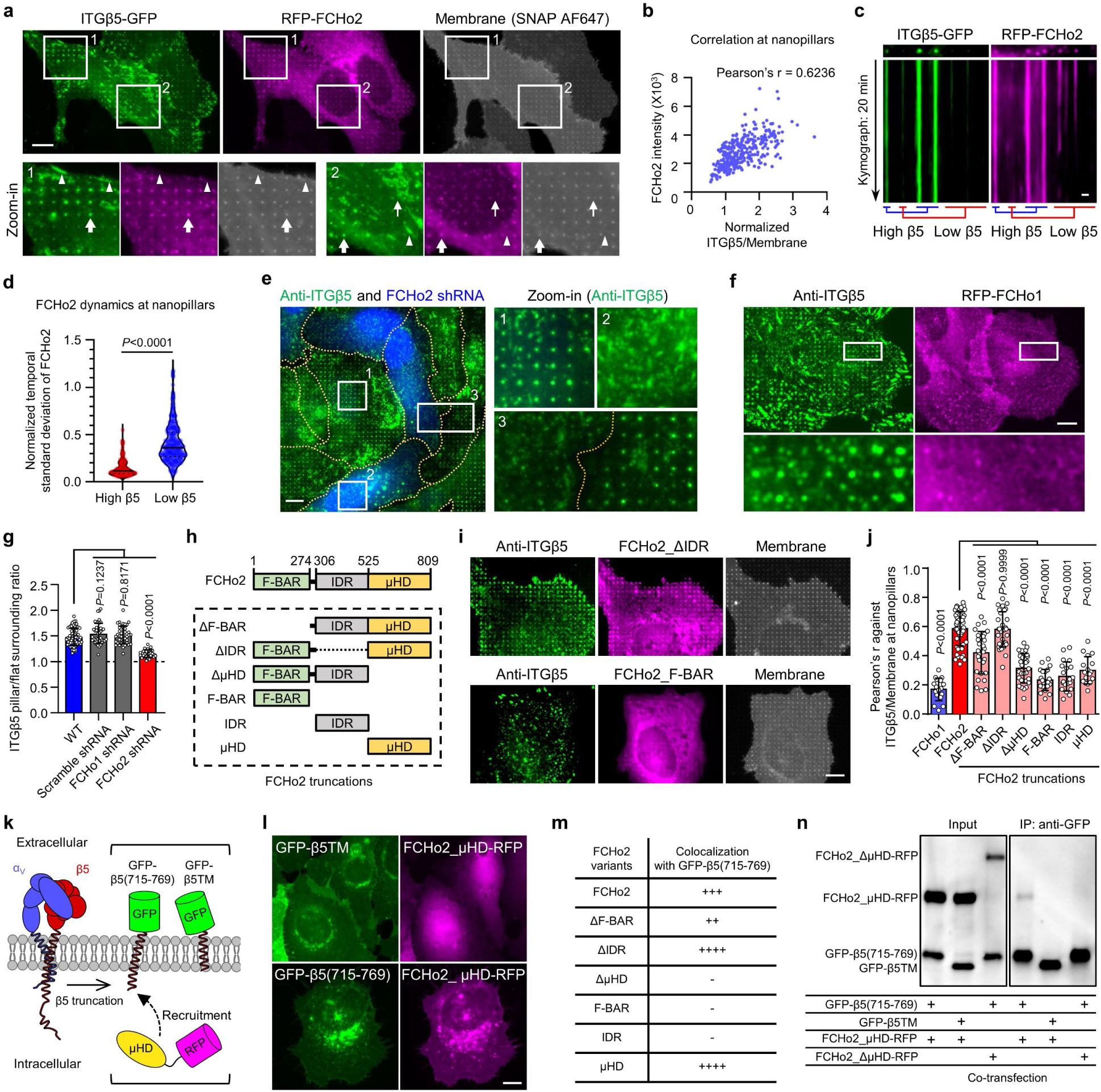
Curved adhesions require a curvature-sensing protein FCHo2. **a**, RFP-FCHo2 positively correlates with ITGβ5-GFP in curved adhesions (arrows) at vitronectin-coated nanopillars. In the same image, FCHo2 is absent in focal adhesions (arrow heads) and clathrin-containing adhesions (thin arrows). **b,** Scatter plot of RFP-FCHo2 intensity against the ITGβ5/membrane ratio at N = 299 nanopillars. **c,** Kymographs of ITGβ5-GFP and RFP-FCHo2 at vitronectin-coated nanopillars imaged at 15 s/frame for 20 min. **d,** The temporal standard deviations of RFP-FCHo2 for 20 min at N = 104 high-β5 and N = 313 low-β5 nanopillars, from three independent cells. Standard deviations values are normalized to the initial intensities. Medians (lines) and quartiles (dotted lines) are shown. **e,** FCHo2 knockdown significantly reduces nanopillar-induced ITGβ5 accumulation. BFP expression is a marker of shRNA transfection. **f,** ITGβ5 does not colocalize with FCHo1 at nanopillars. **g,** Quantification showing that FCHo2 knockdown can significantly reduce nanopillar-induced ITGβ5 accumulation, but FCHo1 knockdown cannot. N = 68/43/44/30 cells. **h,** Illustration of FCHo2 domain organization and truncations. **i,** Representative images showing that FCHo2_ΔIDR correlates with ITGβ5 in curved adhesions at nanopillars, but FCHo2_F-BAR does not. FCHo2_F-BAR overexpression also reduces ITGβ5 accumulation at nanopillars. **j,** Pearson correlation coefficients of FCHo1, FCHo2, and truncated FCHo2 with ITGβ5 at vitronectin-coated nanopillars from N = 17/48/29/29/31/22/20/17 cells. **k,** Illustration of the engineered proteins GFP-β5(715-769) and GFP-β5TM, and the cytosolic protein FCHo2_µHD-RFP. **l,** FCHo2_µHD-RFP is cytosolic and diffusive when co-expressed with GFP-β5TM (top). When co-expressed with GFP-β5(715-769), FCHo2_µHD-RFP localizes to the plasma membrane and colocalizes with GFP-β5(715-769) in the perinuclear Golgi region (bottom). **m,** Qualitative analysis of GFP-β5(715-769)-induced membrane re-localization of FCHo2 variants. **n,** Immunoblots of co-immunoprecipitation assay confirms the interaction between the ITGβ5 juxtamembrane region and FCHo2_µHD. Data are mean ± SD (**g**, **j**), from two independent experiments. *P* values calculated using Mann-Whitney test (**d**) or one-way ANOVA with Bonferroni’s multiple comparison (**g** and **j**). Scale bars: 10 µm (**a**, **e**, **f**, **i**, **l**); 1 µm (**c**).

Interestingly, RFP-FCHo1 does not show strong accumulation at nanopillars (**Fig 4f**). Even when RFP-FCHo1 accumulates on some nanopillars, it shows little correlation with anti-ITGβ5. Furthermore, shRNA knockdown of FCHo1 does not affect ITGβ5 accumulations at nanopillars (**Extended Data Fig. 10c** and quantification in **Fig. 4g**), suggesting that FCHo1 is not involved in curved adhesions. Moreover, membrane wrapping around nanopillars is not affected by knockdown of FCHo1/2 (**Extended Data Fig. 10d**). These results indicate that FCHo2, but not FCHo1, participates in curved adhesions, thus we focus on FCHo2 for further investigation.

FCHo2 is composed of a curvature-sensitive F-BAR domain, an intrinsically disordered region (IDR) that is crucial for activating AP2 in clathrin-mediated endocytosis, and a C-terminal μHD domain^35^ (**Fig. 4h**). We constructed six FCHo2 variants that lack one or two of its three regions. Cells were transfected with each of these FCHo2 variants (GFP-tagged) and immunostained with anti-ITGβ5. Deletion of the IDR does not affect FCHo2_ΔIDR’s colocalization with ITGβ5 in curved adhesions, but all the other FCHo2 variants have either no correlation or reduced correlation with ITGβ5 in curved adhesions (**Fig. 4i,j**). Overexpression of some variants, such as FCHo2_FBAR, induces a dominant-negative effect and a significant reduction of curved adhesions. These results indicate that both the F-BAR domain and the μHD domain, but not the IDR, are necessary for FCHo2’s participation in curved adhesions.

Finally, we investigated whether the cytoplasmic region of ITGβ5 interacts with FCHo2. Based on our ITGβ5 truncation experiments, we identified the juxtamembrane region of the ITGβ5 as a crucial region for curvature sensing. Therefore, we engineered a fusion protein GFP-β5(715-769) that is composed of an extracellular GFP tag and a 55-aa segment ITGβ5(715-769), which contains the transmembrane domain and a short 27-aa intracellular fragment (**Fig. 4k**). GFP-β5(715-769) does not contain ITGβ5’s extracellular domains or its intracellular talin-binding site. As a negative control, we constructed GFP-β5TM, another fusion protein composed of an extracellular GFP and the transmembrane domain of ITGβ5(715-749) but lacking the crucial juxtamembrane fragment (**Fig. 4k**). Both GFP-β5(715-769) and GFP-β5TM are largely localized to the plasma membrane with some accumulations in the perinuclear regions (**Fig. 4l** and **Extended Data Fig. 11**). These perinuclear accumulations are proteins trapped in the Golgi apparatus and colocalize with a Golgi marker Golgi-RFP (**Extended Data Fig. 11a**), which is typical of membrane proteins.

When FCHo2_µHD-RFP is expressed by itself or co-expressed with GFP-β5TM, the protein is highly diffusive in the cytosol, which is expected for the cytosolic µHD domain (**Fig. 4l**, top). However, when FCHo2_µHD-RFP is co-expressed with GFP-β5(715-769), its cellular distribution changes dramatically to be membrane localized (**Fig. 4l**, bottom). FCHo2_µHD-RFP also accumulates and colocalizes with GFP-β5(715-769) in the perinuclear Golgi regions. The redistribution of FCHo2_µHD-RFP suggests its interaction with GFP-β5(715-769). Furthermore, GFP-β5(715-769) co-expression also induces membrane and Golgi localization for the other three µHD domain-containing FCHo2 variants: full-length FCHo2, FCHo2_ΔF-BAR and FCHo2_ΔIDR (**Extended Data Fig. 11b**), but not for the three variants that do not contain the µHD domain: FCHo2_ΔµHD, FCHo2_F-BAR and FCHo2_IDR (**Extended Data Fig. 11c** and **Fig. 4m**). In control experiments where GFP-β5TM is co-expressed, full-length FCHo2, similar to FCHo2_µHD-RFP, does not overlap with GFP-β5TM (**Extended Data Fig. 11d**).

Co-immunoprecipitation experiments further support that the cytoplasmic fragment of β5 interacts with FCHo2_µHD. GFP-β5(715-769) was co-expressed with FCHo2_µHD-RFP in HEK293T cells. When the lysates were incubated with anti-GFP-conjugated beads, FCHo2_µHD-RFP was precipitated together with GFP-β5(715-769) (**Fig. 4n**). However, no FCHo2_µHD-RFP signal was detected when it was co-expressed with GFP-β5TM (**Fig. 4n**). Furthermore, when FCHo2_ΔµHD-RFP, a variant lack the µHD domain, was co-expressed with GFP-β5(715-769), no co-immunoprecipitation signal was detected either (**Fig. 4n**). Thus, our results from both cell-based experiments and in vitro assays indicate a protein-protein interaction between the 20-aa intracellular fragment of ITGβ5 and the µHD domain of FCHo2. This interaction is essential for ITGβ5-mediated curved adhesion.

### Curved adhesions form abundantly on 3D ECM fibres

Based on their distinct molecular compositions, curved adhesions can be identified by the colocalization of ITGβ5 and FCHo2, and focal adhesions by the colocalization of ITGβ5 and vinculin. On flat and rigid glass substrates (**Fig. 5a**), there are many focal adhesions (marked by the overlap of ITGβ5 and vinculin) (**Fig. 5b**), but minimal overlaps between FCHo2 and ITGβ5 (**Fig. 5c**).

**Fig. 5:**
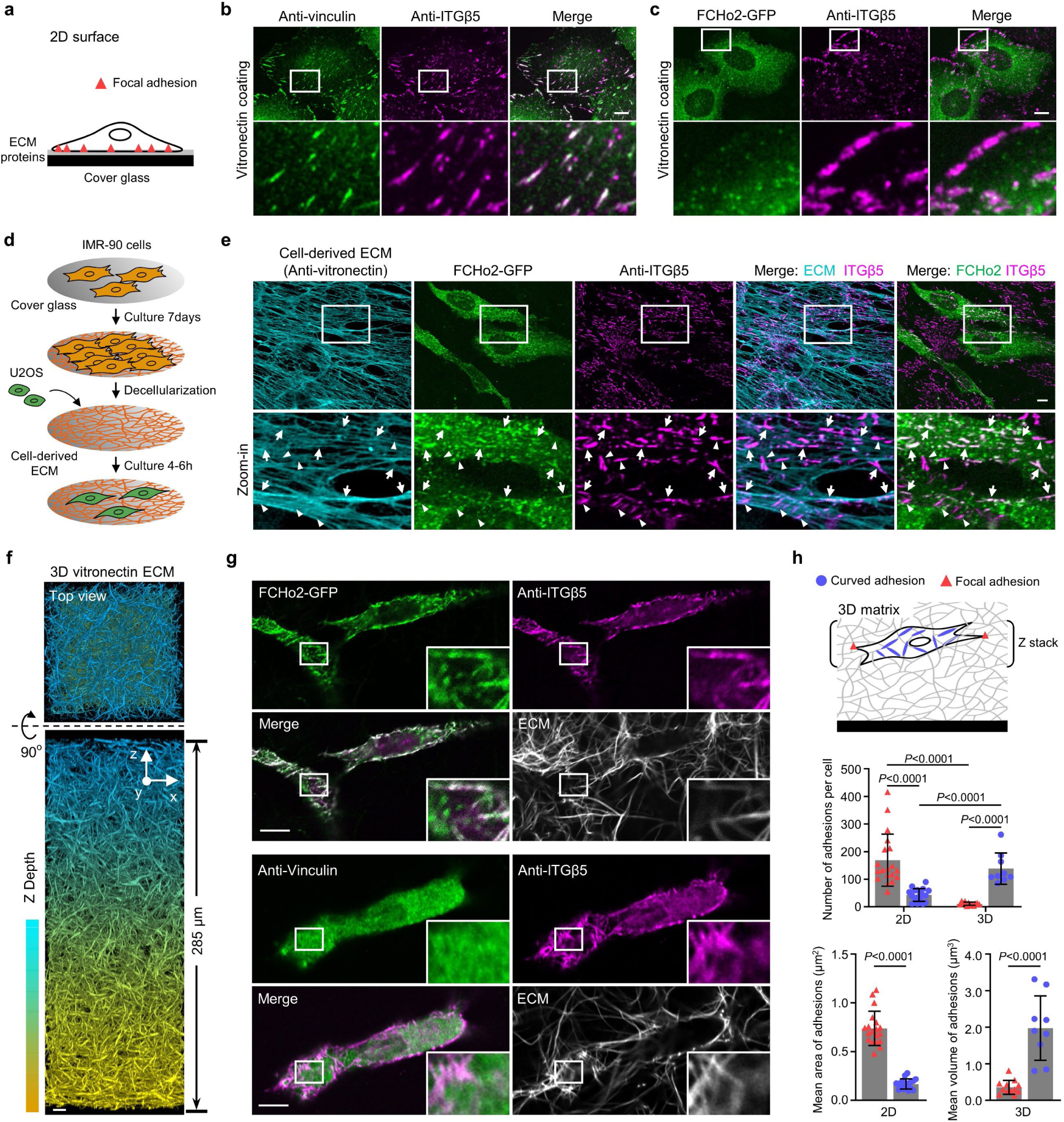
Curved adhesions are prevalent in physiologically relevant environments. **a,** Schematic of a cell growing on ECM-coated 2D flat surfaces. **b,** ITGβ5 and vinculin strongly colocalize on vitronectin-coated flat surfaces. **c,** ITGβ5 and FCHo2 do not colocalize on vitronectin-coated flat surfaces. **d,** Schematic illustrating the generation of cell-derived ECM fibres. **e,** IMR-90-derived ECM fibres have vitronectin incorporated as shown by anti-vitronectin staining. U2OS cells form both curved adhesions (arrows, colocalization of Anti-ITGB5 with FCHo2-GFP) and focal adhesions (arrowheads, Anti-ITGB5 devoid of FCHo2-GFP) on these fibres. **f,** Representative 3D images of a thick layer of matrices made of vitronectin fibres. **g,** Curved adhesions, marked by the colocalizations of ITGβ5 and FCHo2, are abundant on vitronectin fibres in 3D matrices (top), while focal adhesions marked by the colocalization of ITGβ5 and vinculin are sparse (bottom). **h,** Quantifications of the number and size of focal adhesions and curved adhesions on 2D flat surfaces and in 3D matrices of vitronectin fibres in N = 19/18/12/9 cells. Data are mean ± SD, from two independent experiments. *P* values calculated using t-test (**h**-adhesion size) or Kruskal-Wallis test with Dunn’s multiple-comparison (**h**-adhesion number). Scale bars: 10 µm.

The natural ECM is enriched with fibrous structures, which may locally deform the cell membrane to induce the formation of curved adhesions. To probe whether curved adhesions exist in physiologically relevant environments, we first examined whether curved adhesions form on a thin layer of ECM fibres derived by fibroblast cells (**Fig. 5d** and **Extended Data Fig. 12a**). Lung fibroblast IMR-90 cells were cultured for 7 days in the presence of ascorbic acid to enhance the production of collagen-based ECM fibres^36^. After decellularization, we observed a thin (∼ 1 µm thickness) layer of ECM fibres that contain vitronectin as shown by the strong anti-vitronectin staining, which agrees with a previous report^37^ (**Fig. 5e**). U2OS cells expressing FCHo2-GFP were cultured on these cell-derived fibres for 6hrs and then fixed and stained with anti-ITGβ5. Anti-ITGβ5 appears as numerous large and elongated patches resembling adhesion structures. A closer examination shows that there are two distinct populations of ITGβ5 patches: one population shows extensive overlap with FCHo2-GFP (arrows) while the other population is devoid of FCHo2-GFP (arrow heads). As FCHo2 and ITGβ5 only colocalize in curved adhesions but not in focal adhesions, clathrin-containing adhesions, or other known cellular structures, their colocalization indicates that cells form curved adhesions on cell-derived fibres. The population of ITGβ5 patches devoid of FCHo2-GFP are likely focal adhesions. It is interesting to note that curved adhesions always align with underlying fibres, while focal adhesions are often not aligned with and can be perpendicular to the fibre direction (**Fig. 5e**).

To further probe the presence of curved adhesions in 3D, we assembled thick ECM layers made of two different fibres: pure collagen fibres and vitronectin-associated collagen fibres (referred to as vitronectin fibres)^38, 39^. Both types of fibres incorporate 10% of AF647-labelled collagen for visualization. Multimeric vitronectin was used to ensure high affinity vitronectin binding to collagen^40^. Colocalizations of AF647-labeled collagen and anti-vitronectin confirmed the presence of vitronectin in the fibre structures (**Extended Data Fig. 12b**). The most observed diameter of these ECM fibres is ∼115 nm as measured by 3D super-resolution microscopy (**Extended Data Fig. 12c,d**). We used a two-step assembly method to prepare 3D matrices of 250-300 µm thickness (**Methods**, **Fig. 5f** and **Extended Data Fig. 12e**) When U2OS cells expressing GFP-CaaX are plated on top of the 3D ECM matrices, most cells have infiltrated and fully embedded in 3D matrices made of vitronectin fibres after 72 hours of culture, but cells largely stay on the surface of 3D matrices made of pure collagen fibres (**Extended Data Fig. 13a**; **Supplementary Video 3 and 4**). The fibres can clearly deform the plasma membrane (**Extended Data Fig. 13b,c**).

When cells are embedded in 3D vitronectin fibres, anti-ITGβ5 signals in these cells significantly overlap with FCHo2-GFP on vitronectin fibres (**Fig. 5g**, **Extended Data Fig. 14a** and **Supplementary Video 5**), indicating the formation of curved adhesions in 3D matrices. Conversely, anti-ITGβ5 accumulations on ECM fibres rarely overlap with anti-vinculin (**Fig. 5g**, **Extended Data Fig. 14b**, and **Supplementary Video 6**), agreeing with previous reports that focal adhesions are sparse in soft 3D matrices^12, 13^. This contrasts with vitronectin-coated 2D rigid surfaces, where ITGβ5 and FCHo2 do not colocalize while ITGβ5 and vinculin strongly colocalize.

We quantified curved adhesions by the colocalized signals of ITGβ5 and FCHo2, and focal adhesions by the colocalized signals of ITGβ5 and vinculin (See Methods for details). The quantifications further confirm that curved adhesions, not focal adhesions, are the dominant adhesion type in soft 3D environments (**Fig. 5h**).

### Curved adhesions facilitate cell migration in 3D ECMs

Because U2OS cells are able to migrate into 3D ECM made of vitronectin fibres but not 3D ECM made of pure collagen fibres, we hypothesized that the formation of curved adhesions may facilitate cell migrations in 3D matrices. For this study, we examined three commonly used human cancer lines: osteosarcoma U2OS cells, lung carcinoma A549 cells, and cervical carcinoma HeLa cells. These cell lines are not highly aggressive and do not migrate in collagen fibres.

Cells expressing GFP-CaaX and various shRNAs were plated on the top surface of 3D fibrous matrices and cultured for 72 hrs before fixation and imaging with a confocal microscope. **Figure 6a** shows the U2OS cells in 3D matrices. The cells are colour-coded according to their depth in the matrices. On ECM made of vitronectin fibres, wild type cells can migrate >100 µm deep into matrices. To perturb curved adhesions, we used shRNA lentiviral vectors to achieve high knockdown efficiencies of FCHo2, ITGβ5, or both. The disruption of curved adhesion can largely abolish the ability of U2OS cells to migrate into vitronectin ECM. Furthermore, when the 3D matrix is made of pure collagen fibres that do not support curved adhesions, wild-type U2OS cells fail to infiltrate into the matrices.

**Fig. 6:**
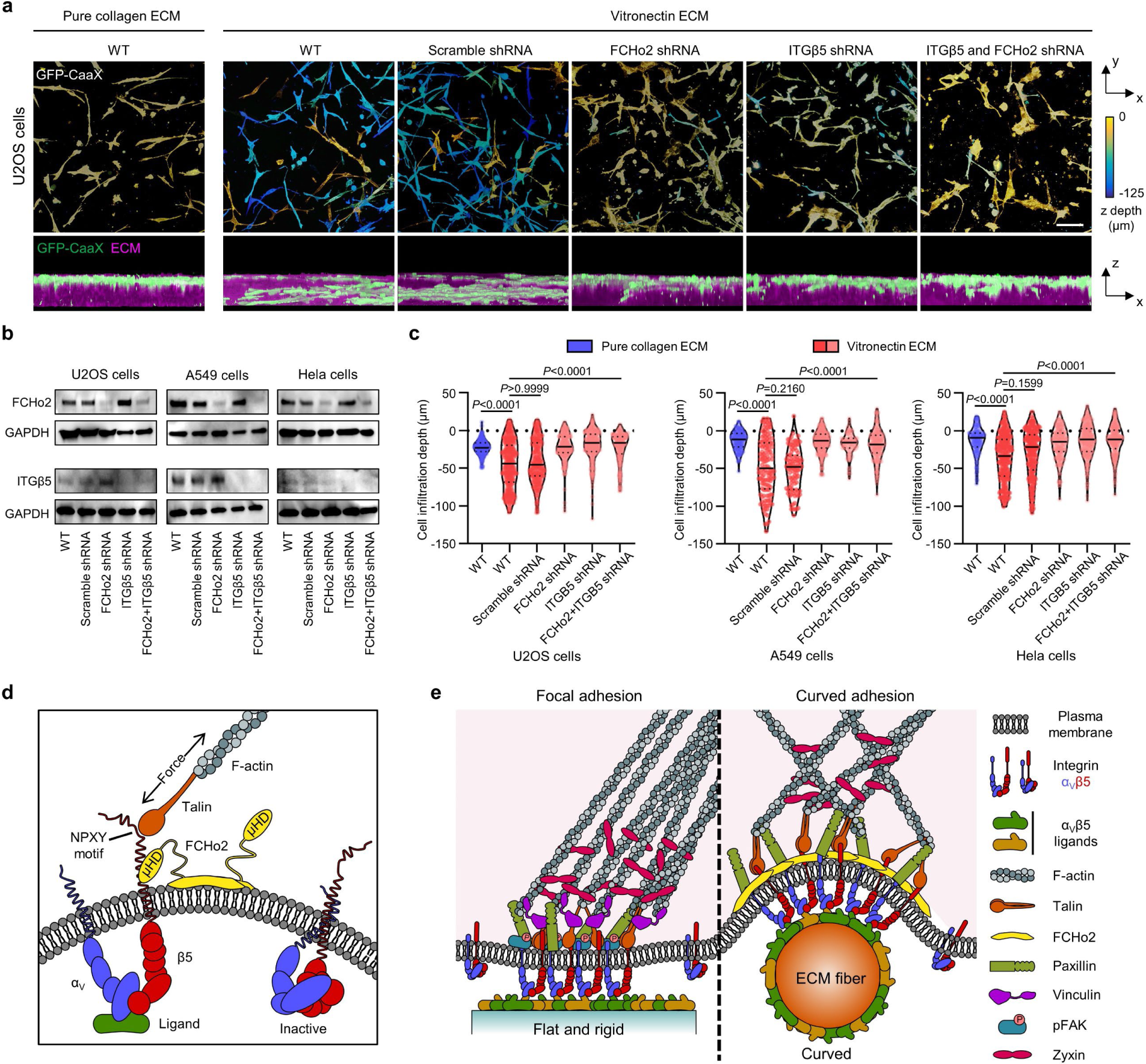
Curved adhesions facilitate the cell migration in 3D ECMs. **a,** Representative images of U2OS cells expressing GFP-CaaX in 3D matrices after 72-hr culture. Cells are colour-coded according to their depth in the matrices in x-y projections. Cells are coloured in green and merged with ECM in magenta in x-z projections. WT cells infiltrated into matrices made of vitronectin fibres, but not matrices made of pure collagen fibres. The shRNAs of FCHo2, ITGβ5, or both, but not scramble shRNA inhibited the cell infiltrations. **b,** Western blots show that shRNAs are able to effectively reduce the expression of FCHo2, ITGβ5, or both in U2OS, A549, and Hela cells. These results support the knock-down efficiencies supplied by the company (Supplementary Table 1). **c,** Quantification of the cell infiltration depth for U2OS, A549, and Hela cells in 3D pure collagen matrices or 3D vitronectin matrices with the knockdown of FCHo2, ITGβ5, or both. N = 100-300 cells for each condition, from two independent experiments. Medians (lines) and quartiles (dotted lines) are shown. *P* values calculated using Kruskal-Wallis test with Dunn’s multiple-comparison. **d,** Illustration of ITGβ5 interacting with FCHo2 in curved adhesions. **e,** Schematic comparison of focal adhesion and curved adhesion architectures. Scale bars: 100 µm.

We find that A549 cells and HeLa cells show similar migration behaviours as U2OS cells (**Extended Data Fig. 15**). Both cell lines are able to infiltrate into vitronectin ECM, but not into pure collagen ECM. HeLa cells infiltrate less deeply than A549 or U2OS cells. Knocking down FCHo2, ITGβ5, or both largely blocks A549 and HeLa infiltration into vitronectin ECM. Western blots show that shRNAs are able to effectively reduce the expression of FCHo2 or ITGβ5 in all three cell lines (**Fig. 6b**), agreeing with the manufacturer’s data of 78-90% knockdown efficiency (**Supplementary Table 1**). Quantifications of a large number of cells (100-300 cells per condition) confirm the observations (**Fig. 6c**), indicating that curved adhesions facilitate cell migration in soft 3D matrices.

## Discussions

Here, we report the identification of curved adhesions, a new type of cell-matrix adhesion that selectively forms on curved membranes and is mediated by the integrin ɑVβ5. Curved adhesions are molecularly and functionally distinct from focal adhesions and clathrin-containing adhesions. Our data supports a model that curved adhesions require an interaction between the juxtamembrane region of ITGβ5 and the µHD domain of FCHo2, while the curvature sensitivity is conveyed through the F-BAR domain of FCHo2 (**Fig. 6d**). Without extracellular ligands, integrin ɑ subunit blocks the access to the intracellular juxtamembrane region of integrin β tail through an ɑ/β salt bridge, which explains why curved adhesions cannot form without extracellular ligands and disassemble upon Ca^2+^ depletion. Membrane curvature likely facilitates the formation of curved adhesions by recruiting FCHo2 that preferentially binds to curved membranes. The self-oligomerization propensity of FCHo2 likely also assists the clustering and activation of ITGβ5 in curved adhesions. This study reveals a new function of FCHo2 in cell adhesions. Curved adhesions engage a unique subset of adhesion proteins, including talin-1, paxillin, and zyxin, but not vinculin or pFAK (**Fig. 6e**). The presence of talin-1 in curved adhesions enables the linkage to the actin cytoskeleton and the transmission of mechanical forces.

An interesting observation is that curved adhesions form in physiologically relevant soft environments and facilitate cell migration in 3D matrices. Both integrin β5 and FCHo2 are widely expressed in tissues^26^ and integrin β5 is upregulated in many types of cancer^41–43^. Vitronectin and fibronectin, the extracellular ligands for curved adhesions, are abundant ECM proteins^44, 45^. The identification of curved adhesions provides new leads for the understanding of complex cell-ECM coupling in biological and pathological conditions.

## Methods

### Antibodies and shRNA oligo sequences

All shRNAs were obtained from Sigma-Aldrich, except for the scramble shRNA which was obtained from Addgene (a gift from David Sabatini, Addgene #1864). The knockdown efficiencies of shRNAs were provided by Sigma-Aldrich using quantitative PCR. To knock down Integrin β5, three different shRNAs, each with an efficiency ∼80%, were combined to further increase the efficiency. For AP2-μ, clathrin heavy chain, FCHo1, FCHo2, EPS15, EPS15R, intersectin 1, and intersectin 2, two different shRNAs were combined to knock down each of these proteins. The clone IDs of shRNA vectors used in this study and their respective knockdown efficiencies are listed in **Supplementary Table 1**. The primary and secondary antibodies used in this study are obtained from commercial sources and listed in **Supplementary Table 2**.

### Nanofabrication of nanobars, nanopillars, and gradient nanobars

The patterns of nanobar, nanopillar, and gradient nanobar arrays were designed using an open-source python package. The fabrication method was adapted from a previous study^23^. Briefly, a 4-inch quartz wafer (Silicon Materials, 04Q 525-25-1F-SO) was diced and cleaned with acetone and isopropanol under sonication. Clean wafers were spin-coated with 275-nm CSAR 6200 (AllResist) and 100-nm Electra 92 (AllResist) e-beam resists. The desired patterns were exposed to e-beam using the JEOL JBX-6300FS system. A 120 nm-thick layer of chromium mask was deposited on the post-exposure chips using the AJA e-beam evaporator and then immediately lifted off with acetone and isopropanol. The Cr mask-patterned chips were then etched anisotropically to create vertical nanostructures by reactive ion etching (Plasma Therm Versaline LL ICP Dielectric Etcher, PT-Ox) with a mixture of C_4_F_8_, H_2_, and Ar for 3 min. Finally, the chips were immersed in chromium etchant 1020 (Transene) for 30 min to remove the Cr mask.

### Fluorescence labelling of gelatin, vitronectin, and collagen

To prepare for gelatin labelling, gelatin (Sigma-Aldrich, G9391) was added into water and autoclaved to make a 2 mg/mL gelatin solution. The gelatin solution was then diluted 2-fold with 0.2 M sodium bicarbonate buffer (Sigma-Aldrich) before labelling. To prepare for vitronectin labelling, the human multimeric vitronectin solution (Molecular Innovations Inc, HVN-U) was adjusted to a concentration of 1 mg/mL and then its buffer was exchanged to 0.1 M sodium bicarbonate using a Zeba spin desalting column (Thermo Scientific, 89882). For fluorescence labelling of gelatin or vitronectin, Atto 647 NHS Ester (Sigma-Aldrich, 07376) or Cy3 Mono NHS Ester (Sigma-Aldrich, GEPA13101) dye was dissolved in DMSO to make 1mg/ml stock solutions and then diluted with 0.1 M sodium bicarbonate solutions to a final concentration of 1 µg/ml. The dye solution was then mixed with the protein solution with a 1:1 molecular ratio to make Atto 647-gelatin or Cy3-vitronectin. The mixtures were incubated for 1 h at room temperature (RT). Then, free dyes were removed, and the buffer was changed to 1x phosphate-buffered saline (PBS, Gibco) using Zeba spin desalting columns.

For collagen labelling, we adapted a protocol from a previous study^39^. Briefly, collagen type I (Corning, 354236) was first diluted into a final concentration of 4 mg/ml with ice-cold 20 mM acetic acid to prevent polymerization. The collagen solution (1 mL) was then neutralized with a pre-chilled mixture of 20 µL 1M NaOH, 200 µL 10x DMEM, 200 µL 10 mM HEPES, and 580 µL water. The neutralized collagen was dropped in a pre-chilled 10-cm petri dish and incubated for 1 h at RT for fibre formation. The collagen fibres were washed with 10 mL PBS 3 times for 10 min each and then washed with 10 mL 0.1 M sodium bicarbonate 3 times for 10 min each to remove non-polymerized collagen. Alexa Fluor 647 (AF647) NHS ester (Thermo Scientific, A37573) was dissolved at 1mg/ml in DMSO and diluted in 0.1 M sodium bicarbonate to a final concentration of 2 µg/mL and then added to the collagen fibres. In this way, Alexa dyes will label collagen at locations that do not interfere with collagen polymerization. After 30-min incubation at RT, the free dye was washed away with PBS 5 times for 15 min each. The labelled collagen fibres were then dissolved in 500 µL of 500 mM acetic acid by overnight incubation at 4°C. The remaining free dye in collagen solution was removed by dialysis against 4 L of 20 mM acetic acid overnight at 4°C. AF647-collagen is usually mixed with unlabelled collagen to make collagen fibres (see fibre preparation section). Samples were stored in dark to avoid photobleaching from the ambient light.

### Surface coating of quartz nanochips with ECM ligands

The nanopillar and nanobar SiO_2_ substrates were first treated with air plasma (Harrick Plasma) for 15 min and then incubated with 0.1 mg/mL PLL (Sigma-Aldrich, P5899) in PBS at 37°C for 1 h. The chips were then washed with PBS 3 times. For PLL surface coating, we stopped here. To coat with other ECM proteins, the nanochips were further incubated with 0.5% (v/v) glutaraldehyde (Sigma-Aldrich, G6257) in PBS at RT for 15 min. After washed with PBS 3 times, the chips were immediately incubated in PBS with the desired ECM protein, such as 1 mg/mL unlabelled gelatin, 1 mg/mL Atto 647-gelatin (described in the fluorescence labelling section), 0.25 mg/ml human plasma fibronectin (Sigma-Aldrich, 341635), 0.25 mg/ml unlabelled vitronectin (PeproTech, 140-09), 0.25 mg/ml Cy3-vitronectin (described in the fluorescence labelling section), 0.25 mg/ml laminin 511 (Sigma-Aldrich, CC160), or 1% (w/v) BSA (Sigma-Aldrich, A9418). After adding the desired protein, the reaction proceeded at 37°C for 1 h and in the dark if the desired protein was fluorescently labelled. The chips were then washed with PBS 3 times. Substrates with fluorescent labelling were immediately imaged to examine the uniformity of surface coating. Before cell seeding, non-fluorescent substrates were treated with 1 mg/ml sodium borohydride (Sigma-Aldrich, 452882) in PBS for 10 min then washed with PBS 5 times to reduce autofluorescence.

### Preparation of fluorescently labelled 3D soft ECMs

To prepare pure collagen fibres, unlabelled and AF647-labelled collagen (described in the fluorescence labelling section) was mixed in a 4:1 (w/w) ratio at a final concentration of 2 mg/ml in 20 mM acetic acid. For collagen polymerization, 50 µL collagen mixture was neutralized with a mixture of 1 µL 1 M NaOH, 10 µL 10x DMEM, 10 µL 10 mM HEPES, and 29 µL water on ice. The neutralized collagen was dropped in Lab-Tek II chambered cover glass (Sigma-Aldrich, Z734853) for 15 µL per well, and incubated at RT for 20 min. The samples were then washed with 20 mM sodium phosphate buffer (pH 7.4) 5 times. This washing step removes free collagen and floating fibres, resulting in a thin layer (∼ 4 µm) of pure collagen fibres immobilized on cover glasses. To generate a thick layer (∼ 280 µm) of pure collagen fibres, neutralized collagen mixture was added to the pre-chilled thin fibre layer for 15 µL per well, and incubated at RT for 45 min. The fibres were then gently washed with 20 mM sodium phosphate buffer (pH 7.4) 5 times. It is important to prepare 3D fibres in two polymerization steps, the first step preparing a thin fibre layer that is attached to the surface and then the second step adding additional monomers where the fibres in the first layer serve as seeds for further polymerization. One-step polymerization usually results in a thin layer regardless of the volume, because most polymerized fibres are not anchored to the surface and are removed during the washing step.

To generate vitronectin fibres, freshly made thick layers of pure collagen fibres described above were incubated in 100 µL of 50 µg/ml multimeric vitronectin (Molecular Innovations Inc, HVN-U) in 20 mM sodium phosphate buffer for 1 h at RT. Multimeric vitronectin is known to bind collagen with high affinity. The resulting fibres were then washed with a 20 mM sodium phosphate buffer 3 times to remove excess vitronectin that is not bound to collagen fibres. Fluorescence imaging of anti-vitronectin and AF647 collagen confirmed the strong binding of vitronectin on collagen fibres.

### Preparation of cell-derived ECM fibres

The protocol is adapted from a previous study^46^. IMR-90 lung fibroblast cells were cultured in complete cell-culture media supplied with with 110 µg/mL sodium pyruvate (Gibco) and 100 μg/mL 2-Phospho-L-ascorbic acid trisodium salt (Sigma-Aldrich, 49752) for 5-7 days to produce cell-derived ECM. Anti-vitronectin staining confirms the presence of vitronectin in these fibres. To remove IMR-90 cells from these fibres, the cultures were incubated for 20 min at 37°C in calcium-and magnesium-free PBS supplied with 5 mM EDTA and 2 M urea for decellularization without cell lysis. The samples were then gently washed with PBS 10 times before seeding new U2OS cells or being used for characterization such as anti-vitronectin staining.

### Plasmids construction

The DNA fragment encoding human integrin β5 was amplified from the pCX-EGFP beta5 integrin receptor (a gift from Raymond Birge, Addgene #14996), and then inserted into the pEGFP-N1 vector (Clontech) for the expression of ITGβ5-GFP. The DNA fragments encoding human integrin β3 and β8 were amplified from the complementary DNA (cDNA) of U2OS cells and were cloned into the pEGFP-N1 vector for the expression of ITGβ3-GFP and ITGβ8-GFP, respectively. The DNA fragment encoding integrin β4 was amplified from pcDNA3.1/Myc-His beta4 (a gift from Filippo Giancotti, Addgene #16039), and then inserted into the pEGFP-N1 vector for the expression of β4-GFP. The expression vector of ITGβ6-GFP was a gift from Dean Sheppard (Addgene #13593).

The expression vector of talin 1-RFP was a gift from Michael Davidson (Addgene #55139). FL TSM and HP35st TSM were gifts from Carsten Grashoff (Addgene plasmids #101170 and #101251). The DNA fragments encoding TSMs were inserted between the head and rod domains of human talin 1 at aa 447 as talin tension sensors or at the N-terminus of human talin 1 as controls. The resulting DNA fragments were used to replace the EGFP fragment in pEGFP-N1 vector for the expression of talin tension sensors and controls. The DNA fragments encoding human zyxin were amplified from the cDNA of U2OS cells and then inserted into the pmCherry-N1 vector (Clontech) for the expression of zyxin-RFP.

The DNA fragments encoding GFP-CaaX and RFP-CaaX were generated by fusing the DNA fragment encoding the CaaX motif of K-Ras protein (GKKKKKKSKTKCVIM) to the 3′-end of the DNA fragments encoding EGFP and FusionRed. The DNA fragment of RFP-CaaX was inserted into the pEGFP-C1 vector (Clontech) to replace the EGFP for the mammal expression. The DNA fragment encoding GFP-CaaX was inserted into pLenti pRRL-SV40(puro)_CMV for lentivirus packaging. The expression vectors of LifeAct-RFP were gifts from Michael Davidson (Addgene #54491). The DNA fragment encoding SNAP-tag was amplified from the pSNAP-tag (m) vector (a gift from New England Biolabs & Ana Egana, Addgene #101135), and then inserted into the pDisplay vector (Invitrogen, V66020) for the expression of cell-surface SNAP-tag.

The shRNAs are cloned in the pLKO.1 vector for transcription in mammalian cells. The pLKO.1 vector is designed for transient transfection and lentivirus transduction. The puromycin resistant sequence in pLKO.1 vector was replaced with a sequence encoding EBFP2 to fluorescently label cells that were transfected or transduced with shRNAs.

The expression vector of RFP-FCHo1, RFP-FCHo2 and EPS15-RFP were gifts from Christien Merrifield (Addgene # 27690, #27686 and #27696). The DNA fragment of FCHo2 is amplified and cloned into a pEGFP-N1 vector to generate FCHo2-GFP. To generate GFP-or RFP-tagged FCHo2 truncations, DNA fragments encoding FCHo2_F-BAR (aa 1-274), FCHo2_ΔF-BAR (aa 275-809), FCHo2_ΔIDR (aa 1-305 fused with aa 520-809), FCHo2_IDR (aa 306-526), FCHo2_µHD (aa 520-809), and FCHo2_ΔµHD (aa 1-525) were amplified and cloned into the pEGFP-N1 or pmCherry-N1 vector.

### Cell lines and cell culture conditions

U2OS (ATCC HTB-96™), A549 (ATCC CCL-185™), HeLa (ATCC CCL-2™), and Mouse Embryonic Fibroblast (MEF) (ATCC CRL-2991™) cells were cultured in complete cell-culture media, Dulbecco’s modified Eagle’s medium (DMEM) (Gibco) supplied with 10% (v/v) fetal bovine serum (FBS) (Sigma-Aldrich), and 1% (v/v) penicillin and streptomycin (Gibco). IMR-90 lung fibroblast cells (ATCC^®^ CCL-186™, a gift from Scott Dixon) were maintained in complete cell-culture media supplied with with 110 µg/mL sodium pyruvate (Gibco) and 100 μg/mL 2-Phospho-L-ascorbic acid trisodium salt (Sigma-Aldrich, 49752) to stimulate the production of cell-derived ECM. HEK293T cells (ATCC^®^ CRL-3216™) were cultured in complete cell-culture media supplemented with 110 µg/mL sodium pyruvate (Gibco) for lentivirus production. Lentivirus were used to achieve high transfection efficiency as well as to generate a stable GFP-CAAX U2OS cell lines. HT-1080 (ATCC CCL-121™), U-251MG (Sigma-Aldrich, 09063001), and MCF7 (ATCC HTB-22™), and human mesenchymal stem cells (PCS-500-011™) were cultured in Eagle’s minimum essential medium (EMEM) (Gibco) supplied with 10% (v/v) FBS, and 1% (v/v) penicillin and streptomycin. All cell lines were maintained at 37°C in a 5% CO_2_ atmosphere.

Before being seeded on nanochips or protein fibres, cells were maintained in a 6-well plate reaching ∼50% confluency. Cells were detached with 10-min incubation in an enzyme-free cell dissociation buffer (Gibco, 13151014). The detached cells were collected by spin at 300 relative centrifugal force (RCF) for 3 min and then resuspended in CO_2_-balanced pre-warmed cell-culture media before cell plating.

### Transient cell transfection

For U2OS, A549, HT1080, U251MG, MCF7, human mesenchymal stem cells, and MEF cells, transient transfections were achieved by electroporation. Cells were grown in 6-well plates at ∼80% confluency. For transfection, one well of cells was detached, collected, and then resuspended in an electroporation buffer containing 100 µL Electroporation buffer II (88 mM KH_2_PO_4_ and 14 mM NaHCO_3_, pH 7.4), 2 µL Electroporation buffer I (360 mM ATP + 600 mM MgCl_2_), and 0.5-2 µg plasmids. The cells were electroporated in 0.2-cm gap electroporation cuvettes by Amaxa Nucleofector II (Lonza) using protocols pre-installed by the manufacturer.

For HeLa cells, transient transfection was achieved by Lipofectamine 2000. HeLa cells at ∼50% confluency in 6-well plates were incubated with 2 mL Opti-MEM-I medium (Gibco, 31985062) supplied with 6 µL Lipofectamine 2000 transfection reagent (Invitrogen) and 1-2 µg plasmids for 2 h. To mark Golgi apparatus, U2OS cells were transfected with N-acetylgalactosaminyltransferase-RFP using CellLight™ Golgi-RFP, BacMam 2.0 virus (Invitrogen, C10593).

For GFP tagged integrin β isoforms (β3-GFP, β4-GFP, β5-GFP, β6-GFP, β8-GFP), transfected cells are cultured for 3 days in a 6-well plate before they are gently detached with 10-min incubation in an enzyme-free cell dissociation buffer (Gibco, 13151014). These detached cells are then replated on ECM-coated substrates and imaged after 6 hrs.

### Lentiviral transduction and stable cell line generation

For lentivirus packaging, HEK293T cells at ∼80% confluency in 35-mm dishes (Corning, 353001) were transfected with 1.5 µg 3rd generation lentivirus package vector encoding GFP-CaaX or shRNAs, 0.8 µg psPAX2 (a gift from D. Trono, Addgene #12260), and 0.7 µg pMD2.G (a gift from D. Trono, Addgene #12259), mixed with 9 µL lipofectamine 2000 transfection reagent in 2 mL Opti-MEM-I medium. The transfection media were replaced with 2 mL of HEK293T culture media 6 h after transfection. The transfected cells were maintained for 24 h. Then, the virus-containing supernatants were collected, and the cell debris was removed by spin at 400 RCF for 5 min followed by filtration through 0.45-μm PVDF syringe filter units (Millipore).

Lentiviral particles were used to transduce U2OS, A549, and HeLa cells with shRNAs to achieve high-efficiency knockdown of ITGβ5, FCHo2 or both. Stable cell lines expressing GFP-CaaX were generated by lentivirus transduction of wild-type U2OS cells followed by antibiotic selection with 2.5 µg/mL puromycin in complete cell-culture media. The antibiotic pressure was released 3 days before experiments.

### Immunofluorescence labelling

The primary and secondary antibodies used in this study are listed in **Supplementary Table 2**. For the immunolabeling of integrin β1 or αvβ5 with primary antibodies that recognize extracellular domains, cells were first snap-chilled in ice-cold HEPES-buffered hanks balanced salt solution (HBSS), which is 1X HBSS (Gibco, 24020117) buffered with 10 mM HEPES (Gibco, 15630106). The samples were then incubated with primary antibodies, which were 1:100 diluted in HHBSS, for 30 min at 4°C. The samples were then washed with ice-cold HHBSS 3 times for 5 min each and immersed with 4% ice-cold PFA in PBS to fix for 15 min at RT. If talin co-staining was required afterward, the samples were fixed with 4% PFA in a PHEM buffer (PIPES 60 mM, HEPES 25 mM, EGTA 10 mM, MgCl_2_ 2 mM, pH 6.9) instead of 4% PFA in PBS. The fixed samples were washed and permeabilized with 0.1% Triton-X (Sigma-Aldrich, T8787) in PBS for 15 min at RT. Then, the permeabilized samples were washed and incubated in a blocking buffer (5% BSA in PBS) for 1 h at RT. The blocked samples were incubated in primary antibodies 1:250 diluted in the blocking buffer for 1 h at 37°C and then 1 h at RT.

For the immunolabeling of ITGβ5 (with antibody recognizing ITGβ5’s intracellular fragment), vinculin, paxillin, pFAK (Tyr397), AP2-α, clathrin heavy chain or vitronectin, samples were fixed with 4% PFA in PBS for 15 min at RT. If talin co-staining was required afterward, the samples were fixed with 4% PFA in the PHEM buffer instead. The samples were then washed and permeabilized with 0.1% Triton-X in PBS for 15 min at RT. The permeabilized samples were washed and incubated in the blocking buffer for 1 h at RT. The blocked samples were incubated in primary antibodies 1:250 diluted in the blocking buffer for 1 h at 37°C and then 1 h at RT.

The primary antibody-labelled samples were then washed with the blocking buffer 3 times for 15 min each and incubated in fluorescently labelled secondary antibodies 1:500 diluted in the blocking buffer for 1 h at RT. The samples were washed with the blocking buffer 3 times for 15 min each and then with PBS 3 times before imaging. For the immunostaining of cells embedded in 3D fibres, the time durations for antibody incubation and washing were doubled.

SNAP-tag labelling for membrane visualization (Fig. 3b, 4a; Extended Data Fig. 6) In order to preserve the green and red channels for protein labelling in triple-channel imaging experiments, we avoided using GFP-CaaX or RFP-CaaX. Instead, we used extracellular SNAP-tag to label the cell membrane with Fluor 647. Briefly, cells were transfected with SNAP-pDisplay (the transmembrane domain of PDGFRβ fused with an extracellular SNAP tag). The cell-surface SNAP-tags were fluorescently labelled before fixation. Briefly, cells were incubated in CO_2_-balanced pre-warmed media supplied with 1 µM O^6^-benzylguanine (BG)-coupled Alexa Fluor 647 (NEB, S9136S) for 15 min at 37°C in a 5% CO_2_ atmosphere. The cells were then washed 5 times with cell culture media before fixation.

#### Western blotting of integrin β5 and FCHo2 in U2OS, A549, and HeLa cells

Lentiviral infections were used to transduce cells with shRNAs targeting scramble control, integrin β5, FCHo2, or both. The same procedures were used for U2OS, A549, and HeLa cells. After 72-hr culture, lentiviral infected cells were rinsed with ice-cold 1X PBS and lysed for 1 hr at 4°C in 1X RIPA buffer (25 mM Tris–HCl, 150 mM NaCl, 1% Triton X-100, 1% sodium deoxycholate, 0.1% sodium dodecyl sulfate) supplemented with protease and phosphatase inhibitor cocktails (Roche, 04693159001 and 04906837001). For probing endogenous ITGβ5, the samples were further boiled for 10 min at 95°C. The samples were then spun for 40 min at 4°C after lysis to clarify lysates. The lysates were mixed with 2X Laemmli sample buffer (Bio-Rad, 1610747) and β-mercaptoethanol, boiled for 5 min at 95°C, and subsequently subject to SDS-PAGE using the Mini-PROTEAN Vertical Electrophoresis system (Bio-Rad, 1658026FC). After electrophoresis, the samples were transferred onto a nitrocellulose membrane using the Trans-Blot Turbo Transfer System (Bio-Rad, 1704150). The membranes were blocked with 3% BSA in 1X TBST buffer and then incubated with anti-integrin β5, anti-FCHo2, or anti-GAPDH overnight at 4°C. The protein bands were visualized using HRP-conjugated secondary antibody and chemiluminescence under Azure Imaging Systems (Azure biosystem).

### Co-Immunoprecipitation (Co-IP) assay

HEK293T cells (ATCC^®^ CRL-3216™) were transiently co-transfected with GFP-ITGβ5(715-769) or GFP-ITGβ5TM together with mCherry-FCHo2_µHD or mCherry-FCHo2_ΔµHD by electroporation. After 48-hr culture, cells were washed with ice-cold 1X PBS followed by exposure to freshly prepared 0.5 mM DSP (3,3’-Dithiobis(succinimidyl Propionate), Sigma-Aldrich) in 1X PBS for 30 min at room temperature. Cells were then incubated in an ice-cold 50 mM Tris-HCl buffer for 15 min and lysed in 1X RIPA buffer for 1 hr at 4°C. The lysates were incubated with equilibrated GFP-Trap magnetic agarose (ChromoTek) overnight at 4°C. The beads were pelleted and washed with 1X RIPA buffer for 4 times. The pellets were subsequently resuspended in 2X Laemmli sample buffer (with β-mercaptoethanol) and boiled for 10 min at 95°C to elute and denature proteins. The samples were subject to SDS-PAGE and Western blotting. The blots were incubated with anti-GFP and anti-mCherry overnight at 4°C, and the protein bands were visualized using HRP-conjugated secondary antibodies and by chemiluminescence.

### Epi and Confocal fluorescence microscopy

Fluorescence images were acquired in an epi-fluorescence microscope (Leica DMI 6000B) using a 100x (1.40 NA), 60x (1.40 NA) or 20x (0.80 NA) objective. The microscope is equipped with ORCA-Flash4.0 Digital CMOS camera, Lumencor SOLA light source, and filter sets: 370-39/409/448-63 nm (blue emission), 484-25/505/524-32 nm (green emission), 560-32/581/607-40 nm (red emission), and 640-19/655/680-30 nm (far-red emission). During live-cell imaging, cells were maintained in phenol red-free DMEM (Gibco) supplied with 10% FBS at 37°C with 5% of CO_2_ in a stage top incubator (Tokai Hit, INUBSF-ZILCS). Time-lapse images were taken at 15 s/frame using 500 ms exposure times. Confocal images were acquired in a Nikon A1R confocal microscope using a 20X objective (0.75 NA) or a 40X water immersion objective (WD = 0.61 mm, 1.15 NA). The microscope is equipped with 405, 488, 561, and 633 nm lasers for excitation. Z-stack images were taken every 0.5 µm with 1.2 Airy Unit. Ratiometric FRET imaging of living cells expressing talin tension sensors was performed using an epi-fluorescence microscope (Leica DMI 8000) with excitation at 496-516 nm from Leica LED8 system. The donor emission at 527– 551 nm and the acceptor emission at 602–680 nm were recorded simultaneously using K8 Scientific CMOS microscope camera.

### Expansion microscopy imaging of the cell-nanopillar interface (Extended Data Fig. 8a)

Expansion microscopy was carried out as previously described^32^. Briefly, following cell culture on nanopillar chips, fixation, and immunostaining, samples were incubated overnight at RT in a 1:100 dilution of AcX (Acryloyl-X, SE, 6-((acryloyl)amino) hexanoic acid succinimidyl ester, Invitrogen A20770) in PBS. After washing with PBS, cells were then incubated for 15 min in the gelation solution (19% (w/w) sodium acrylate, 10% (w/w) acrylamide, 0.1% (w/w) N,N’-methylenebisacrylamide) at RT. Then, nanochips were flipped cell side down onto a 70 μL drop of the gelation solution supplemented with 0.5% N,N,N’,N’-tetramethylenediamine and 0.5% ammonium persulfate on parafilm and incubated at 37 °C for 1 h. After gelation, the nanochip with the hydrogel still attached was incubated in a 1:100 dilution of Proteinase K in digestion buffer (50 mM Tris HCl (pH 8), 1 mM EDTA, 0.5% Triton X-100, 1 M NaCl) for 7 h at 37 °C. Hydrogels were then soaked twice in MilliQ water for 30 min and then incubated overnight in MilliQ water at 4 °C. To image samples, excess water on the hydrogels was carefully removed using a Kimwipe. Then, the hydrogels were mounted onto PLL-coated glass coverslips to prevent sliding during imaging.

### Super-resolution microscopy (Extended Data Fig. 12d,e)

Our method is adapted from protocols described previously^47, 48^. The 3D single-molecule data was acquired using a custom-built wide-field double-helix point-spread function (PSF) inverted microscope. The samples were incubated in a blinking buffer solution containing 10% (w/v) glucose (BD Difco), 100 mM tri(hydroxymethyl)aminomethane-HCl (Thermo Fisher), 2 μL/mL catalase, 560 μg/mL glucose oxidase, and 10 mM of cysteamine (all Sigma Aldrich), allowing for a low emitter concentration during data acquisition. The focus was first set to the coverslip using a bead as a reference. Standard localization-based approaches fitted the shape of the PSF of emitters to a 2D Gaussian to yield highly precise XY positions. To extract the Z position, a double-helix phase mask was inserted in the Fourier plane of the microscope to modify the shape of the standard PSF to now have two lobes that rotate as a function of Z. We fitted the double-helix PSF to two Gaussian functions and extracted the midpoint to estimate the XY position. From the fit, we determined the lobe angle of the double-helix PSF. Then, using a carefully calibrated curve that relates lobe angle to Z position, we determined the Z position of the emitter. The localization precision was calculated from the detected photons using a formula calibrated specifically for our microscope. After processing, poorly localized emitters (XY precision > 20 nm or Z precision > 40 nm or lobe distance > 8 pixels) were removed from the reconstruction. Localizations were merged to correct for over counting before quantification of any individual fibre diameters. The localized single-molecule positions were rendered with the Vutara SRX program (Bruker).

### Scanning electron microscopy

Nanostructures without cells were imaged by SEM (FEI Nova NanoSEM 450 and FEI Magellan 400 XHR). For SEM imaging of cells, the cells on nanostructures were washed with PBS and fixed in 0.1 M sodium cacodylate buffer (pH 7.2) supplemented with 2% glutaraldehyde (Electron Microscopy Sciences) and 4% paraformaldehyde (PFA) for 20 min at RT and then overnight at 4°C. The cells were then incubated in 0.1 M sodium cacodylate buffer supplied with 1% OsO_4_ and 0.8% potassium ferricyanide (Electron Microscopy Sciences), for 2 h at 4°C. The samples were dehydrated sequentially with 50%, 70%, 95%, 100%, and 100% (v/v) ethanol in water for 10 min each, and then dried by critical point drying. Before being imaged by SEM, the cell samples were sputter-coated with 3 nm-thick Cr.

## Quantitative analysis methods

All images were prepared and processed using ImageJ (Fiji). Quantifications were performed using ImageJ and MATLAB 2017a (MathWorks).

### Quantification of nanostructure sizes (Extended Data Fig. 1)

The following parameters were measured in Fiji by counting pixels: the height, width, and length of bars; the height, diameters (top, middle, and bottom) of nanopillars. If the samples were tilted 45°, a correction factor was applied to the Z dimension. The diameters of curvatures at bar ends were measured using the Kappa curvature analysis plugin (open source) in Fiji.

### Quantification of the nanobar end/side ratio (Fig. 1i, 1k, 3d, 3i; Extended Data Fig. 9c, 10b)

Since nanobars are regularly spaced, the locations of nanobars were automatically detected from the bright field images using a MATLAB program, which generates an array of evenly spaced square masks with nanobars at the centre. These square masks were averaged to generate a smooth and averaged image of nanobar. This averaged image was then used to identify the bar-end and the bar-side locations with respect to the averaged square mask. Then, these relative locations were translated to individual square masks to identify the bar-end and the bar-side locations for individual nanobars using Fiji. These ROIs were then applied to the integrin or membrane colour channels.

For each nanobar, the end/side ratio of ITGβ5 was calculated and then normalized by dividing with the end/side ratio of CaaX, except in Fig. 3i where the membrane is not labelled and the end/side ratio of ITGβ5 was directly calculated. The end/side ratios of individual nanobars in a cell were averaged and reported as a single data point. For cells on gradient bars, the normalized end/side ratios of the same size nanobars in each imaging field were averaged and reported as a single data point. This is because very few bars with the same diameter were covered by one single cell. Step-by-step description of the process with examples was described in detail in our recent work (Suppl. Fig. 2 in Ref. ^49^).

### Quantification of the ITGβ5/membrane ratios on nanopillars (Fig. 2d, 2g; Extended Data Fig. 4c)

Nanopillars are also regularly spaced like nanobars. The square masks with nanopillars at the center were generated by using the same custom MATLAB code as described for nanobars. For nanopillars, we use CaaX membrane images instead of bright-field images to generate the square masks as the CaaX images had better qualities. Based on the averaged square mask of the plasma membrane channel within each cell, the ROIs for “at nanopillar” and “flat region surrounding nanopillar” were respectively defined as 0-9 and 10-20 pixel from the center of a nanopillar. These ROIs were then applied to each mask region in both integrin and membrane channels. Mean intensities in each ROI were measured. The ITGβ5/membrane ratio for each ROI was calculated and normalized by the mean of ITGβ5/membrane ratio averaged over the entire cell area and reported as a single data point. The normalized ITGβ5/membrane ratios of individual ROIs were reported as mean ± SD.

### Quantifications of FRET ratios (Fig. 2i)

The method was adapted from previous studies^28, 29^. The fluorescent images of donor (Ypet) were auto-thresholded using the ‘Moments’ method in Fiji to generate binary maps for all adhesions. The bright-field images and the custom MATLAB code described above were used to generate binary location maps for nanopillars, in which the ROI of a nanopillar is defined as 0-9 pixels to the nanopillar centre. The binary location map of nanopillars was multiplied by that of all adhesions to generate the mask for curved adhesions. The rest adhesion ROIs are defined as focal adhesions (clathrin-containing adhesions do not contain talin-1). The averaged ratiometric FRET values (acceptor emission intensity/donor emission intensity) in curved adhesions and focal adhesions were calculated for each cell and reported as a single data point.

### Calculation of cell areas (Fig. 2m)

The background of all images was subtracted using a rolling ball (radius, 1000 pixel) tool in Fiji. For cell area measurements, the membrane signal was auto-thresholded using the ‘Li’ method in Fiji to identify cell bodies which were then selected as ROIs. The number of pixels in these ROIs were measured as cell areas and reported as mean ± SD.

### Pearson correlation coefficient at nanopillars (Fig. 3b, 4b, 4j)

To analyze the correlation between a protein of interest (POI) and the ITGβ5/membrane ratio at nanopillars, the location masks for nanopillars were first generated as described above. The means of POI, ITGβ5 and membrane (Cell-surface SNAP-tag AF647) at each nanopillar (0-9 pixel to the nanopillar centre) were measured. For each cell, the ITGβ5/membrane ratios at individual nanopillars were calculated and normalized by its median value in the cell. From two arrays of POI intensities and ITGβ5/membrane ratios at each nanopillar, their Pearson correlation coefficient in each cell was calculated using Prism 9.

### Dynamic measurements of F-actin and FCHo2 fluctuations at nanopillars (Fig. 4d; Extended Data Fig. 7e)

The location masks for nanopillars were generated as described above. For cells on gelatin, ITGβ5 appears uniform on the cell membrane and across all nanopillars. The intensities of LifeAct at all nanopillars (0-9 pixel to the nanopillar centres) in a time-lapse image stack were measured. For cells on vitronectin, the intensities of ITGβ5 at individual nanopillars were first quantified. For F-actin dynamic measurements, the nanopillars with top 25% and bottom 25% in ITGβ5 intensities were grouped as high-β5 nanopillars and low-β5 nanopillars, respectively. For FCHo2 dynamic measurements, the nanopillars with top 25% and bottom 75% in ITGβ5 were separated into two groups: high-β5 nanopillars and low-β5 nanopillars, respectively. The FCHo2 intensities in time series were normalized by their values at time 0. The temporal standard deviations of LifeAct or normalized FCHo2 intensities at individual nanopillars were reported as mean ± SD for each group.

### Quantification of fibre diameters (Extended Data Fig. 12c,d)

Individual collagen fibres were isolated by the 3D super-resolution reconstruction. Multiple transverse line profiles that are 100-nm thick were drawn along each of the isolated fibres. The 3D localizations were projected along these lines into 30-nm bin widths. The histogram was fitted to a Gaussian, and the full width half maximum was extracted to estimate the fibre diameter.

### Quantification of curved adhesions and focal adhesions for 2D flat surfaces and 3D ECM (Fig. 5h)

For quantifications of the number and the area of curved adhesions and focal adhesions in 2D, two-channel fluorescence images (anti-ITGβ5 and anti-vinculin, or anti-ITGβ5 and FCHo2-GFP) were separately auto-thresholded using the ‘Moments’ method in Fiji (no custom input parameters). This step removes diffusive background and identifies protein accumulations. The resulting binary image of ITGβ5 was multiplied by the binary image of FCHo2 to generate overlapping ROIs for curved adhesions. Similarly, the ITGβ5 binary image was multiplied by the vinculin binary image to generate overlapping ROIs for curved adhesions. To eliminate noises and artifacts, the overlapping ROIs were then analyzed with a low threshold of 10 pixels in Fiji using the analyze-particle tool. Then, the number and the average size (area) of remaining ROIs for each cell was automatically quantified using the analyze-particle tool. The analysis is automatically processed with the same threshold and is carried out to ensure unbiased comparison between curved adhesion and focal adhesions. Therefore, the analysis classifies occasional ITGβ5-FCHo2 colocalizations on flat surfaces as curved adhesions even though they are likely random events and not real curved adhesions.

For quantifications of the number and the volume of curved adhesions and focal adhesions in 3D, two-channel confocal 3D-stack images were auto-thresholded to remove diffusive background and segmented using the 3D interactive thresholding segmentation tool of the 3Dsuit plug-in in Fiji. No custom input parameters were needed. This step is to identify protein accumulations. The resulting binary 3D stack of ITGβ5 was multiplied by the binary 3D stack of FCHo2 to generate 3D overlapping ROIs for the quantification of curved adhesions. Similarly, the binary 3D stack of ITGβ5 was multiplied by the binary 3D stack of vinculin to generate 3D overlapping ROIs for focal adhesions. Subsequently, the number and the average volume of adhesions for each cell was measured using the 3D objects counter tool in Fiji with a low threshold of 25 voxel. Similar to 2D analysis, the analysis was carried out with the same parameter applied for both sets of data to avoid bias.

### Quantification of the cell depth in 3D matrix (Fig. 6c)

Confocal 3D-stack images of GFP-CaaX were used to identify cell positions. The first step involves the application of the 3D edge filter of the 3Dsuit plug-in in Fiji to the GFP-CaaX 3D stack to identify the cell boundaries. The second step involves segmenting the 3D cell boundary stack using FIJI Macro 3D ART VeSElecT to identify 3D ROIs for individual cells. For this step, a low volume threshold of 500 voxel was applied to remove cell debris and extracellular vesicles. The third step calculates the centre coordinates (x, y, z) of each cell ROIs using 3D objects counter tool in Fiji.

To determine the depth for each cell, we first calculated the surface height position. Confocal 3D-stack images of AF647 labelled-collagen were subject to surface peeler Fiji macro to identify the surface of the 3D matrix. In this process, a gaussian blurring filter (radius 50 pixel) was applied to fill the gaps between fibres. This step identifies a z-height value for each (x, y) position. Then, the depth of a cell is calculated by subtracting the z-position of the cell centre with the z-position of the surface at the (x, y) location of cell centres. All cells in 3D stacks were pooled together to generate the cell depth distribution.

## Statistics and reproducibility

All data were displayed and statistically analyzed by using Prism 9 (GraphPad software). All experiments were repeated independently at least two times to ensure reproducibility. The normal distribution of each sample group was verified using the D’Agostino-Pearson normality test (α = 0.05). For parametric datasets, we used unpaired two-tailed Welch’s t-test to compare two groups, and one-way ANOVA with Tukey’s or Bonferroni’s multiple-comparison to compare more than two groups. For nonparametric datasets, we used Mann-Whitney test to compare two groups, and Kruskal–Wallis test with Dunn’s multiple-comparison to compare more than two groups. The exact *P* values and sample sizes (N) are denoted in the figures and figure legends. Significance was considered at *P* < 0.05. The test statistics for null hypothesis testing including confidence levels and degrees of freedom are listed in **Supplementary Table 3**.

## Acknowledgements

This work was supported by NIH grants 1R35GM141598 and 1R01GM128142, Ono Pharma Breakthrough Initiative Award, and a Packard Fellowship for Science and Engineering. The nanofabrication in this work was performed at Stanford Nanofabrication Facility under the NSF National Nanotechnology Coordinated Infrastructure program and Stanford Nano Shared Facilities supported by the National Science Foundation under award ECCS-2026822. Authors were also supported by Stanford University Center for Molecular Analysis and Design fellowship (to C-H.L.), National Institutes of Health grant Biotechnology Training Grant fellowship (to M.L.N.), and Molecular Biophysics Training Program T32 GM136568 (to C.E.L.). We thank Taylor Jones for insightful discussions, Dr. Scott Dixon for the lung fibroblast cells, Dr. W.E. Moerner for generous assistance with 3D super-resolution imaging, and Dr. Carolyn Bertozzi for the confocal microscope.

## Author contributions

W.Z. and B.C. conceived the study and designed the experiments. C-T.T., X.L. and Z.J. fabricated nanostructures, and performed SEM measurements. W.Z., C-H.L., Y.Y., C.E.L. and M.L.N. constructed plasmids. Y.Y. and B.C. developed MATLAB codes for data analysis. W.Z., C-H.L. and C.E.L. performed most cell experiments and fluorescent microscopy. M.L.N. performed expansion microscopy. W.Z., C-H.L. and C-T.T. analysed most data. A.R.R. carried out super-solution imaging, reconstruction, and analysis. W.Z., C-H.L., M.L.N., C.E.L., and B.C. wrote the manuscript. All authors discussed the results and contributed to the manuscript.

## Ethics declarations

Competing interests: The authors declare no competing interests.

## Extended Data Figures

**Extended Data Fig. 1:**
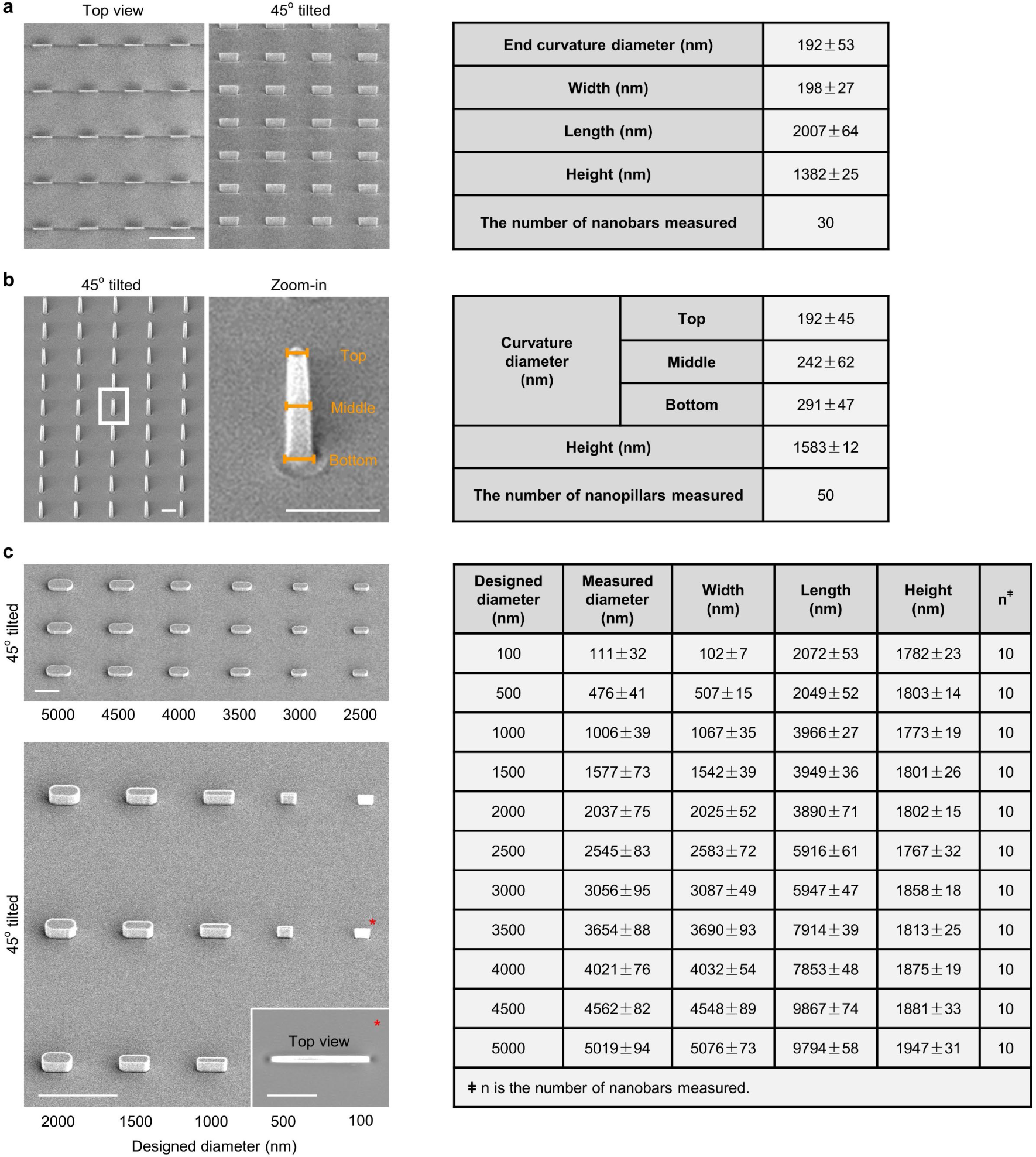
Characterizations of nanobars, nanopillars and gradient nanobars. **a,** Representative SEM images of 200 nm-wide vertical nanobar array. The same sample was either viewed from the top (left) or at 45° angle (right). Scale bar: 5 µm. Statistical analysis of the geometrical dimensions for nanobars is shown in the table. Data are mean ± SD. **b,** Representative SEM images of a vertical nanopillar array. The sample was tilted 45°. Zoom-in image shows one single nanopillar. The top, middle, and bottom diameters of each nanopillar were measured. Scale bars: 1 µm. Statistical analysis of the geometrical dimensions for nanopillars is shown in the table. Data are mean ± SD. **c,** Representative SEM images of gradient bar array with designed end curvature diameters that range from 100 to 5000 nm. The sample was tilted 45°. Scale bars: 10 µm. Inset: the top view of a 100 nm-wide nanobar. Scale bar, 1 µm. Statistical analysis of the geometrical dimensions for gradient bars is shown in the table. Data are mean ± SD.

**Extended Data Fig. 2:**
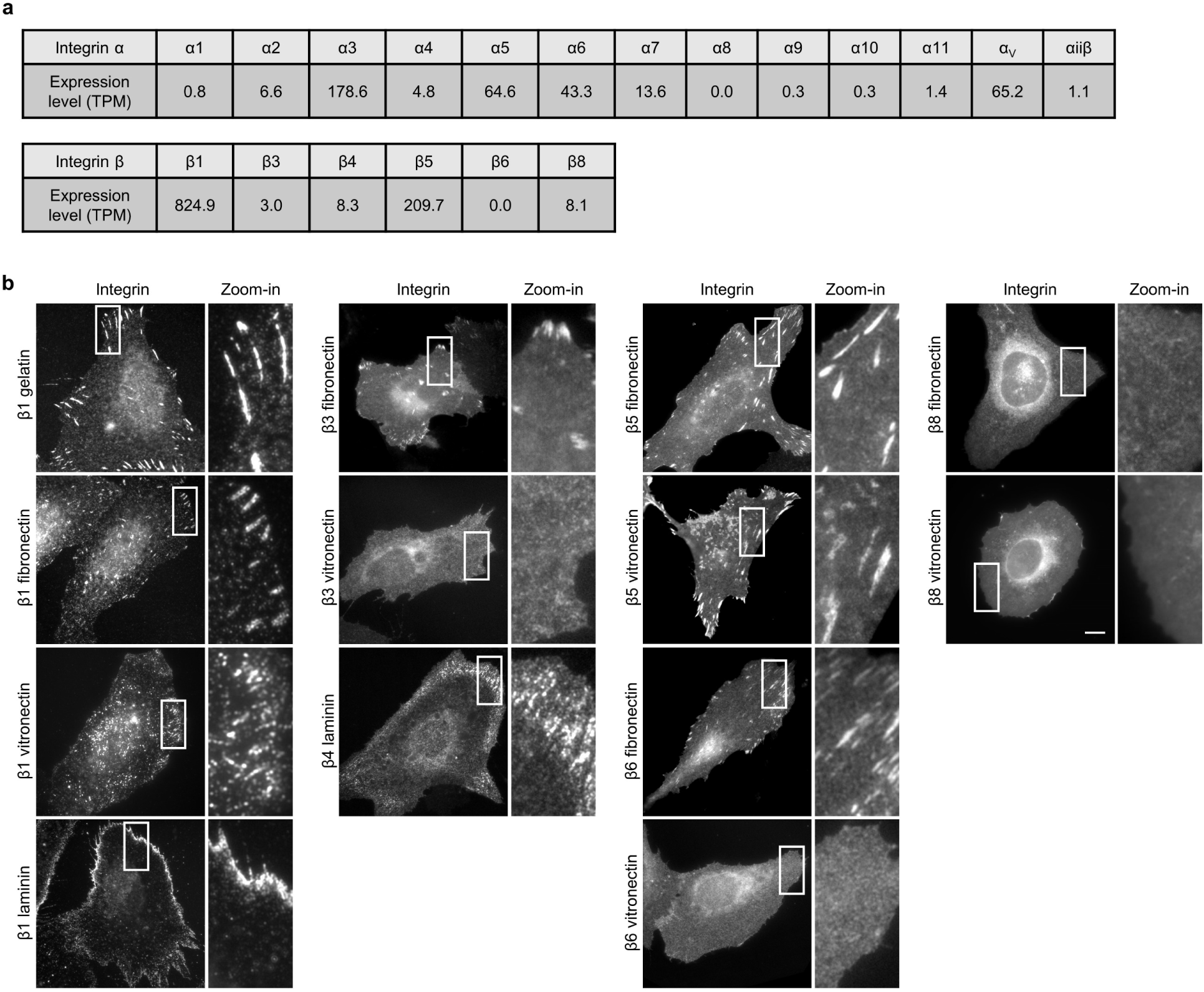
On flat surfaces, integrins form focal adhesions on their respective ECM ligands. **a,** The mRNA levels of the integrin ɑ and β subunits (presented in Fig. 1d) in U2OS cells are listed according to the human protein atlas and a previous study^26^. The mRNA levels are shown in the unit of transcripts per million (TPM). **b,** Fluorescence images of integrin β subunits on flat surfaces coated with different ECM proteins (gelatin, laminin, fibronectin or vitronectin). Endogenous β1 was immunolabeled; β3, β4, β5, β6, and β8 were tagged with GFP and transiently expressed. Scale bar: 10 µm.

**Extended Data Fig. 3:**
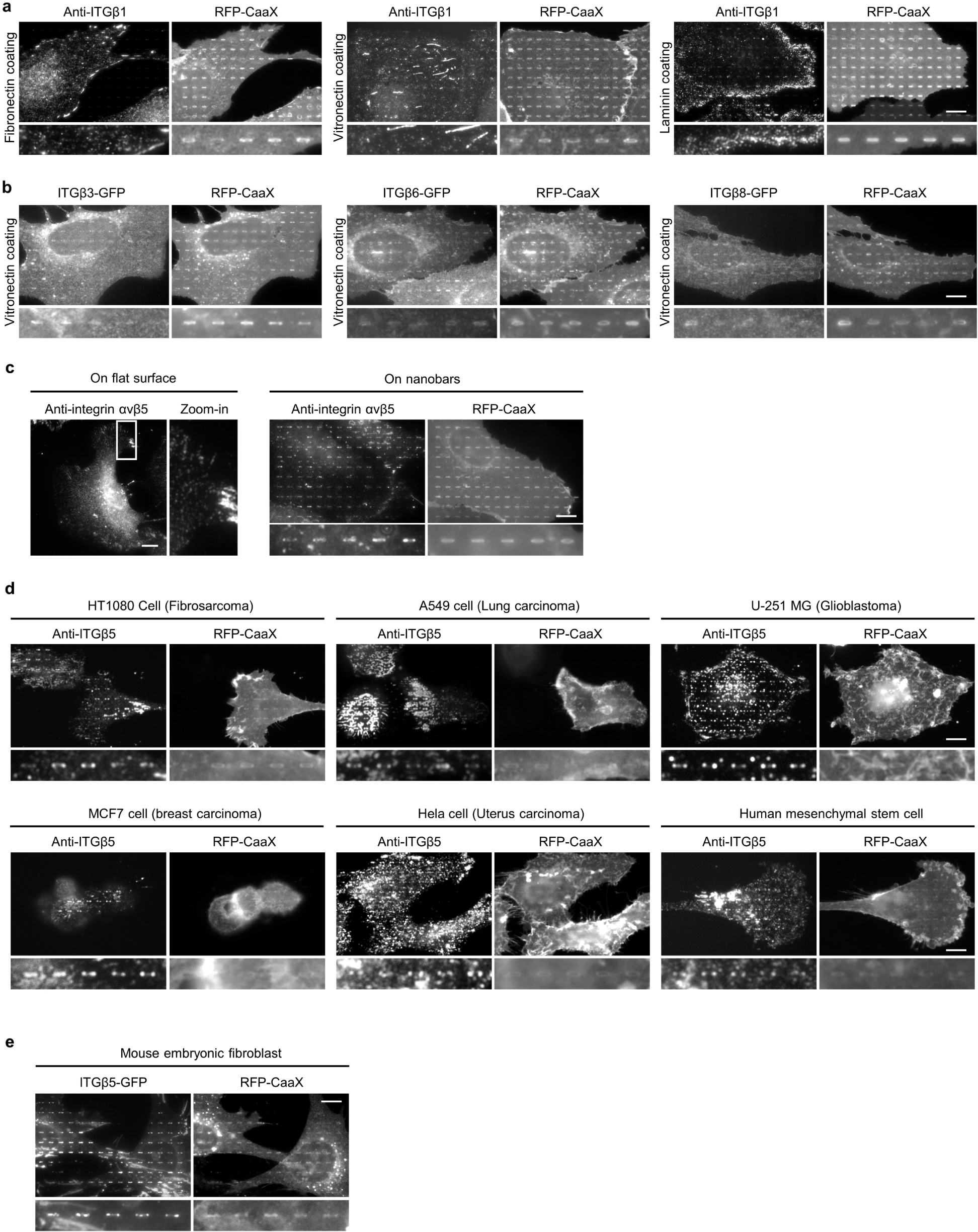
Positive membrane curvature induces the preferential accumulation of integrin ɑVβ5 but not other integrin β isoforms. **a** and **b,** Fluorescence images of endogenous integrin β1 (**a**) and transiently expressed GFP-tagged integrin β3, β6, and β8 (**b**) in U2OS cells expressing a plasma membrane marker RFP-CaaX on 200-nm nanobar arrays coated with different ECM proteins (fibronectin, vitronectin or laminin). All of them show no preference for the nanobar ends. Quantifications of their curvature preferences are presented in Fig. 1i. **c,** Fluorescence images showing that anti-ɑVβ5 in U2OS cells appears in focal adhesions on vitronectin-coated flat surfaces (left), and preferentially accumulates at the ends of vitronectin-coated nanobars in U2OS cells expressing RFP-CaaX (right). **d,** Fluorescence images showing that anti-ITGβ5 preferentially accumulates at the ends of vitronectin-coated nanobars in HT1080, A549, U-251 MG, MCF7, HeLa, and human mesenchymal stem cells. **e,** Fluorescence images showing that ITGβ5-GFP preferentially accumulates at the ends of vitronectin-coated nanobars in mouse embryonic stem cells. Scale bars: 10 µm.

**Extended Data Fig. 4:**
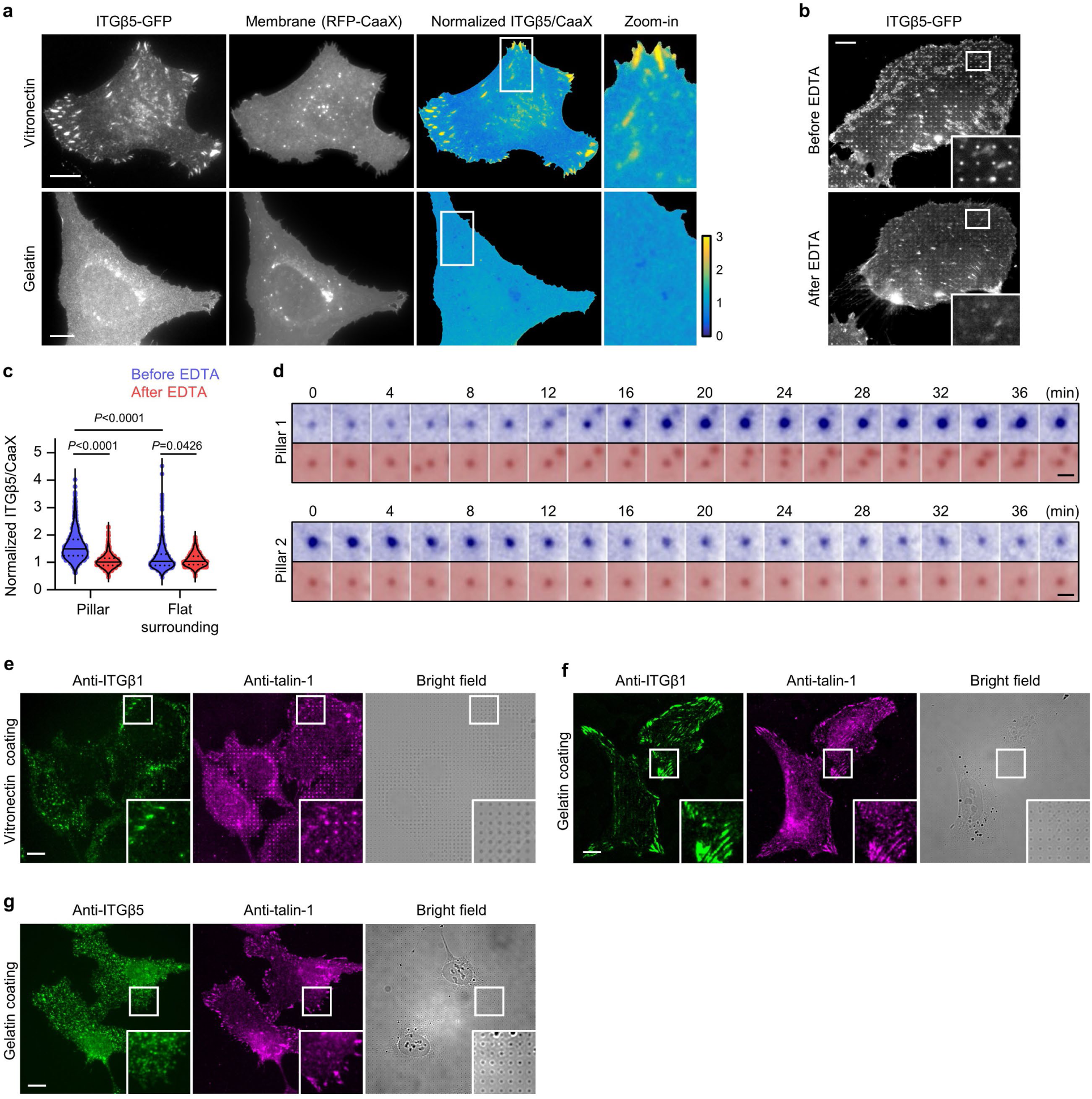
Integrin β5 preferentially accumulates at nanopillars in a ligand dependent manner and recruits talin-1. **a,** Fluorescence images of ITGβ5-GFP and plasma membrane marker RFP-CaaX on vitronectin-coated (top) and gelatin-coated (bottom) flat surfaces. Ratiometric images of ITGβ5/membrane (normalized by its mean per cell) are shown in the Parula colour scale. Scale bars: 10 µm. **b,** Ethylenediaminetetraacetic acid (EDTA) treatment induces a dramatic reduction of ITGβ5-GFP accumulation at nanopillars in a live U2OS cell. Scale bar: 10 µm. **c,** Quantification of the normalized β5/membrane ratio at nanopillars and their surrounding flat regions. N = 968/691 nanopillars before/after EDTA. Medians (lines) and quartiles (dotted lines) are shown. *P* values calculated using paired Kruskal-Wallis test with Dunn’s multiple-comparison. **d,** Time-lapsed images of ITGβ5-GFP and CellMask Orange at Pillar 1 and 2 in Fig. 2e. At Pillar 1, the curved adhesion gradually assembled. At Pillar 2, the curved adhesion gradually disassembled. Scale bars: 1 µm. **e,** Fluorescence and bright-field images showing that ITGβ1 does not accumulate at vitronectin-coated nanopillars, while talin-1 does. **f** and **g**, Fluorescence and bright-field images of immunolabeled talin-1 together with ITGβ1 (**f**) or ITGβ5 (**g**) on gelatin-coated nanopillar substrates. Talin-1 colocalizes with ITGβ1 in focal adhesions, but does not accumulate at nanopillars. ITGβ5 appears mostly diffusive on gelatin-coated substrates. These results indicate that nanopillar-induced talin accumulation depends on ITGBβ5 and its ligands. Scale bars: 10 µm.

**Extended Data Fig. 5:**
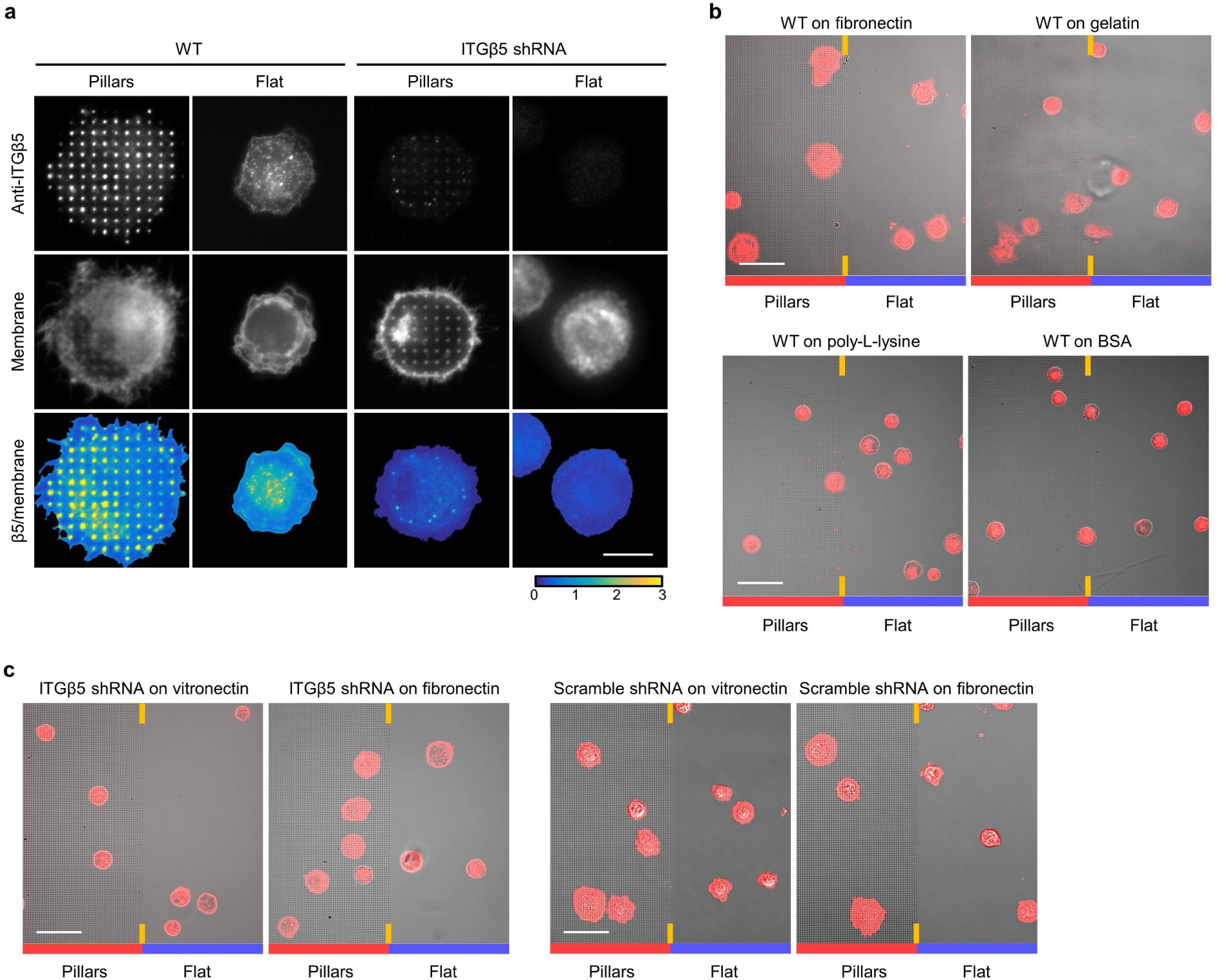
Curved adhesions promote early-stage cell spreading. **a,** ITGβ5 accumulates at vitronectin-coated nanopillars within 30 min after seeding. Cells spread to a larger area on nanopillars than that on flat areas in the same culture. ITGβ5 knockdown (KD) with shRNAs largely abolished the nanopillar-induced early cell spreading. Scale bar: 10µm. **b**, Overlay of the bright-field and fluorescence images of wild type (WT) U2OS cells expressing GFP-CaaX on both nanopillars (left half) and flat surfaces (right half) in the same imaging fields. The cells were seeded on substrates coated with fibronectin, gelatin, poly-L-lysine, or BSA and cultured for 30 min before fixation. Scale bar: 50 µm. **c,** Overlay of bright-field and fluorescence images of U2OS cells transduced with ITGβ5 and scramble shRNAs and stably expressing GFP-CaaX on both nanopillar arrays (left half) and flat surfaces (right half) in the same images. The cells were seeded on substrates coated with vitronectin or fibronectin and cultured for 30 min before imaging. Scale bars: 50 µm.

**Extended Data Fig. 6:**
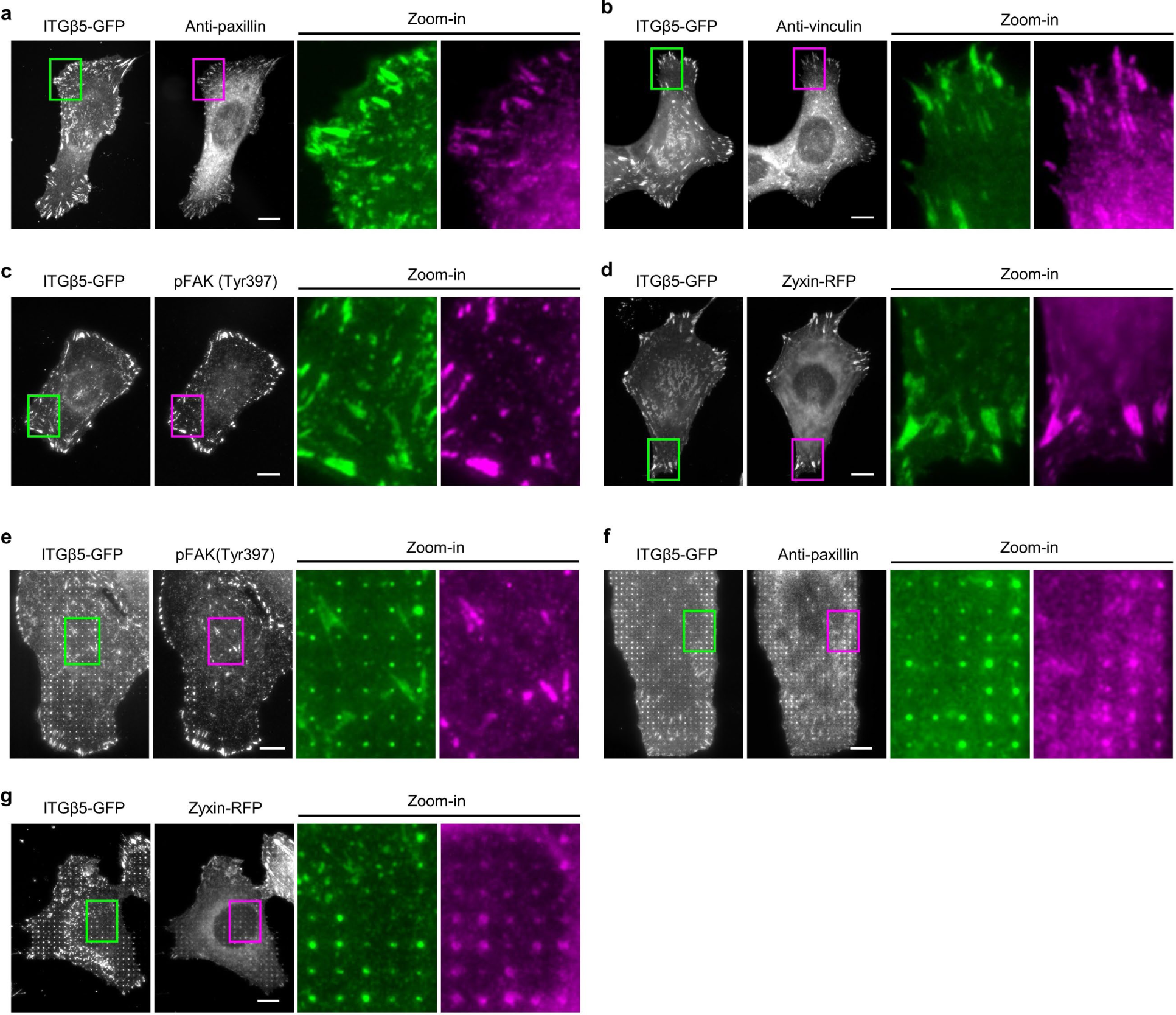
ITGβ5 colocalizes with focal adhesion proteins on flat surfaces, but only a subset of them in curved adhesions. **a** to **d**, Fluorescence images showing that ITGβ5-GFP colocalizes with focal adhesion proteins paxillin (**a**), vinculin (**b**), pFAK (Tyr397) (**c**), and zyxin (**d**) at focal adhesion-like patches in U2OS cells on vitronectin-coated flat surfaces. ITGβ5 shows strong colocalizations with paxillin, vinculin, pFAK, and zyxin in focal adhesions. **e** to **g**, Representative fluorescence images showing that ITGβ5-GFP accumulation at vitronectin-coated nanopillars spatially correlates with paxillin (**f**) and zyxin (**g**), but not with pFAK (Tyr397) (**e**) in U2OS cells. The cell membrane was marked with transiently expressed surface SNAP-tag conjugated with AF647. The normalized ITGβ5/membrane was used to measure ITGβ5 accumulation for the correlation quantification in Fig. 3b. Endogenous paxillin, vinculin, and pFAK (Tyr397) were immunolabeled. Zyxin was tagged with RFP and transiently expressed. Scale bars: 10 µm. The zoom-in images of ITGβ5 (green) and focal adhesion proteins (magenta) are shown.

**Extended Data Fig. 7:**
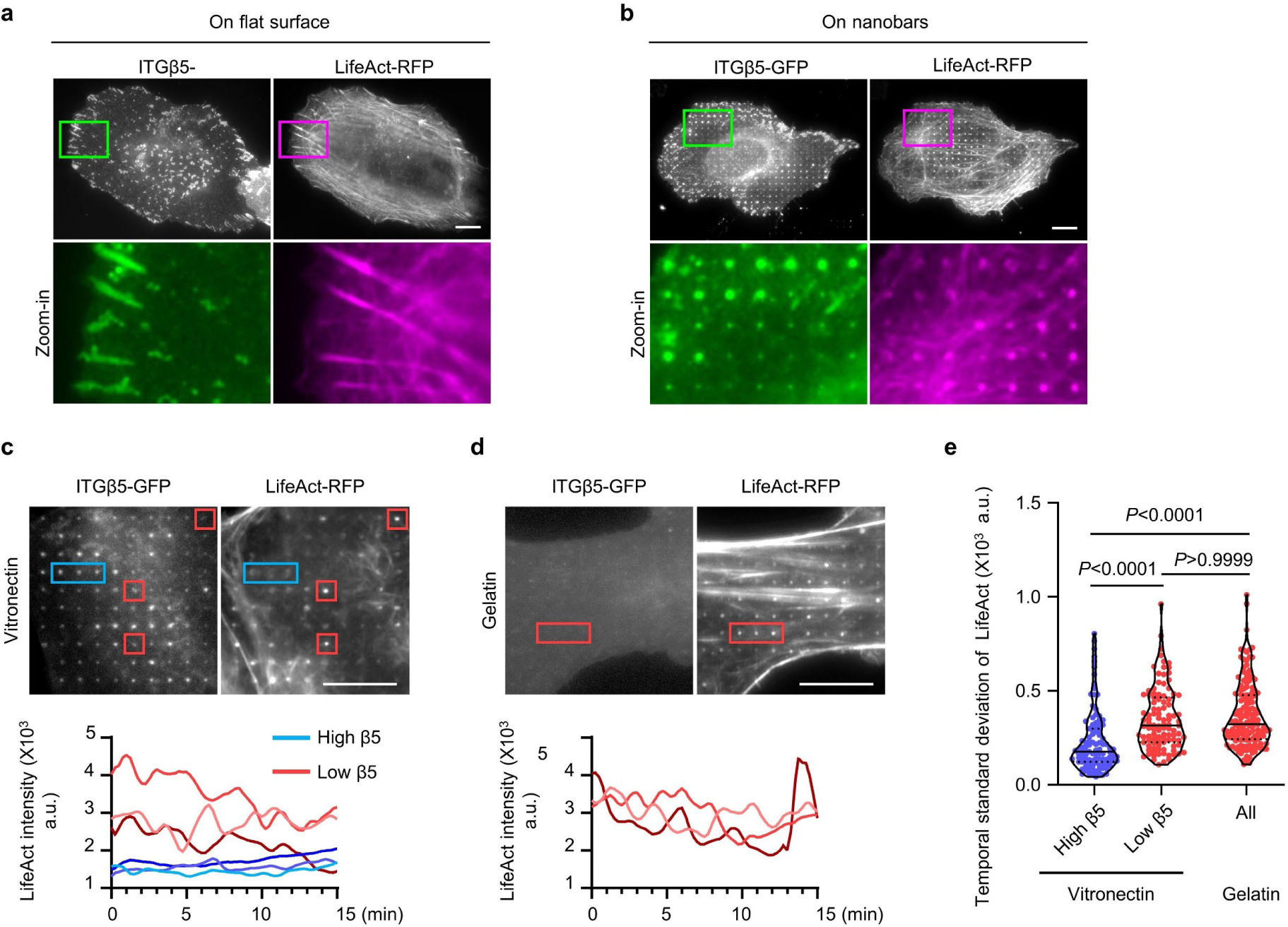
Curved adhesions are linked to a stable population of actin filaments at nanopillars. **a,** On flat areas, actin stress fibres (F-actin labeled by LifeAct-RFP) are anchored to ITGβ5-marked focal adhesion patches. **b,** On nanopillar areas, both ITGβ5 and F-actin preferentially accumulate at nanopillars. However, nanopillars with high β5 accumulations usually do not have high levels of F-actin. A previous study shows that the membrane curvature around gelatin-coated nanopillars is sufficient to induce actin accumulation, but the accumulation is highly dynamic with a lifetime of 1-2 minutes^50^. As curved adhesions are stable and do not form on gelatin-coated nanopillars, we hypothesize that curved adhesions involve a different population of F-actin. **c,** Dynamic correlation between ITGβ5-GFP and LifeAct-RFP in cells cultured on vitronectin-coated nanopillars. There are two distinct populations of the F-actin that accumulate at nanopillars: a stable population of F-actin at high-β5 nanopillars that does not show significant fluctuations, and a dynamic population F-actin at low-β5 nanopillars that rapidly assembles and disassembles within 1-2 minutes. The F-actin intensity of the dynamic population is usually brighter than that of the stable F-actin population. **d,** On gelatin-coated nanopillars that do not induce β5 accumulation, only the highly dynamic population of F-actin is observed at nanopillars. **e,** Quantifications of time-dependent fluctuation confirm two F-actin populations: a stable population on high-β5 nanopillars and a dynamic population at low-β5 nanopillars. The dynamic population of F-actin on vitronectin-coated nanopillars is similar to the dynamic F-actin on gelatin-coated nanopillars. Therefore, curved adhesions involve the stable subpopulation of F-actin at nanopillars. N = 103 high-β5, N=103 low-β5 vitronectin-coated nanopillars, and N = 153 gelatin-coated nanopillars (gelatin coated). Medians (lines) and quartiles (dotted lines) are shown. *P* values calculated using Kruskal-Wallis test with Dunn’s multiple-comparison. Scale bars: 10 µm.

**Extended Data Fig. 8:**
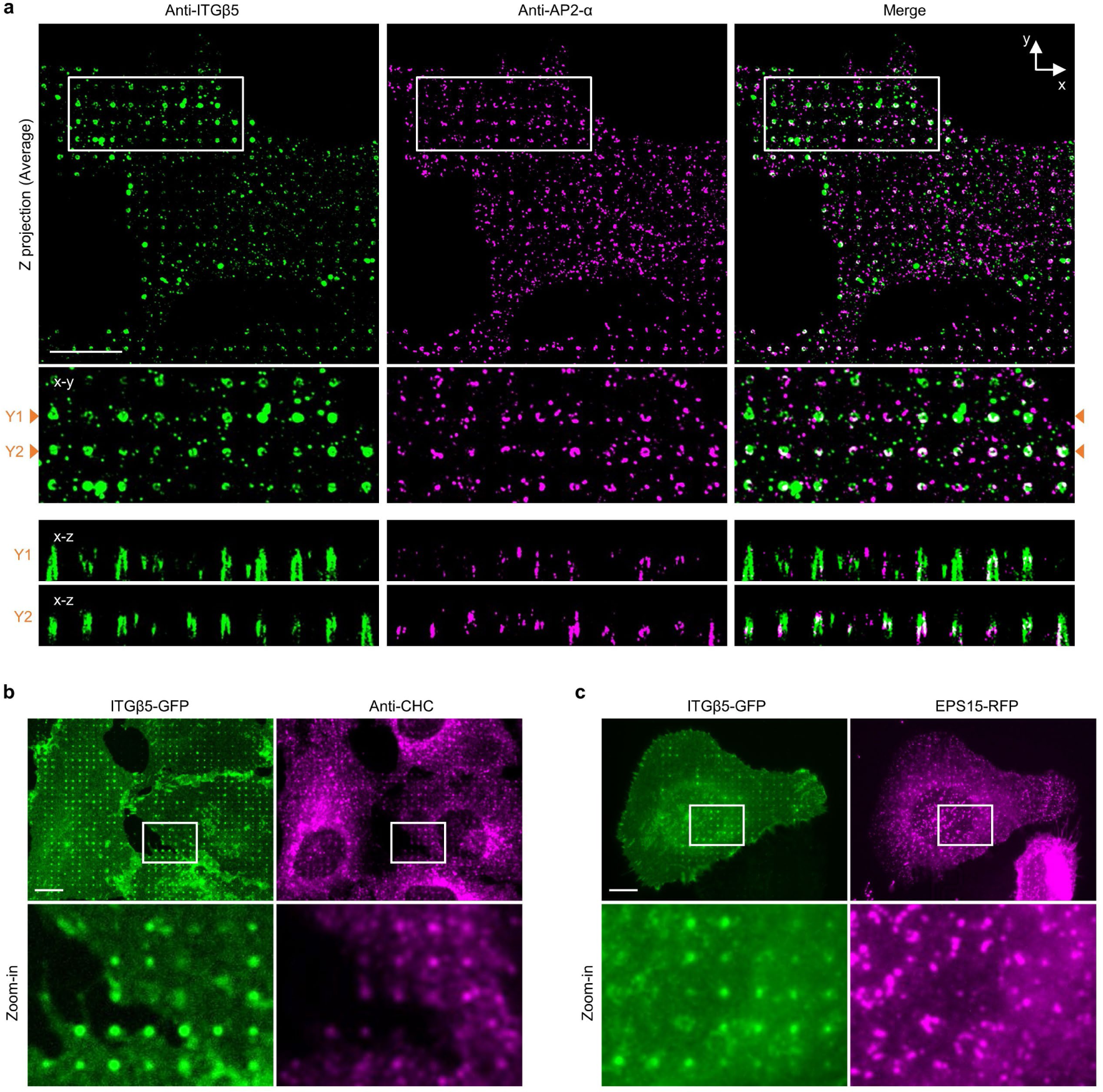
Expansion microscopy shows that integrin β5 and AP2 are not spatially correlated at the nanopillar-membrane interface, and the integrin β5 is not spatially correlated with clathrin and EPS15 at nanopillars. **a,** Expansion microscopy is used to increase the spatial resolution of optical imaging for the nanopillar-membrane interface (see the method section and Ref. ^32^ for detailed descriptions). Both ITGβ5 and AP2-α are immunolabeled. Top row: x-y images of the Z projection (Average) and zoom-in images of the area indicated by white boxes. Bottom: x-z images showing the distribution of immunolabeled ITGβ5 and AP2-α along nanopillars at y = Y1 and y = Y2 in the zoom-in images. Even when ITGβ5 and AP2-α accumulate on the same nanopillar in the x-y image, they are not correlated in the z-dimension. **b**, Both ITGβ5-GFP and immunolabeled clathrin heavy chain (CHC) accumulate at vitronectin-coated nanopillars, but their intensities are not correlated, i.e. nanopillars with high intensities of ITGβ5-GFP are usually not the nanopillars with high intensities of anti-CHC. **c**, ITGβ5-GFP accumulates at vitronectin-coated nanopillars, but the co-transfected EPS15-RFP does not show strong accumulation or correlation with ITGβ5-GFP at these nanopillars. Scale bars: 10 μm.

**Extended Data Fig. 9:**
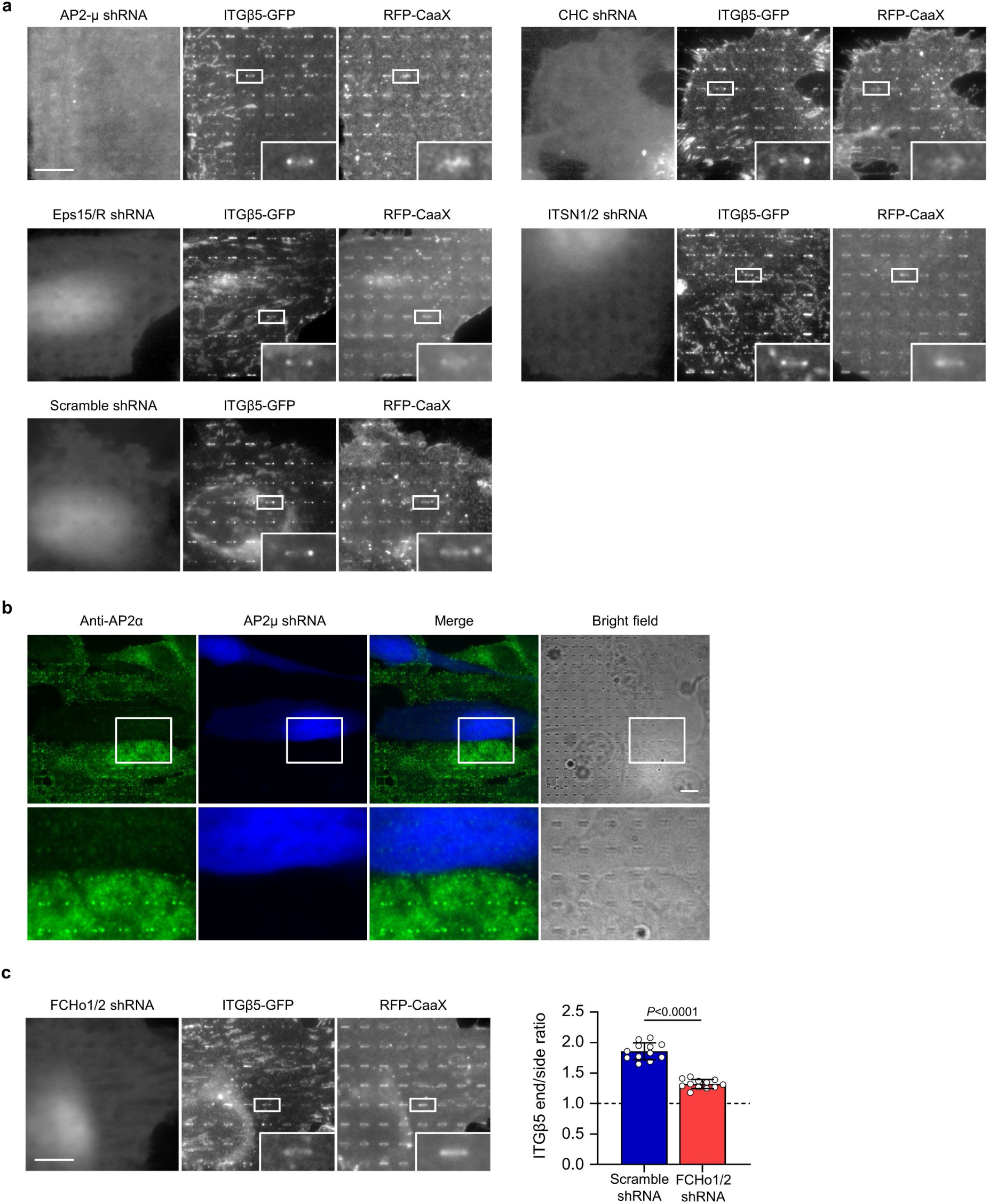
The curvature preference of integrin β5 is not reduced upon the knockdown of AP2-μ, clathrin heavy chain (CHC), EPS15/R, Intersectin1/2, but is significantly reduced upon the knockdown of FCHo1/2. **a,** Representative fluorescence images showing that the shRNA knockdown of AP2-μ, clathrin heavy chain (CHC), EPS15/R, or Intersectin1/2 does not affect the accumulation of ITGβ5-GFP accumulation at the ends of vitronectin-coated nanobars in U2OS cells expressing RFP-CaaX membrane marker. BFP expression is a marker of shRNA transfection. Quantifications of the β5-GFP curvature preference under these conditions are presented in Fig. 3d. **b,** Immunofluorescence showing that the transfection of AP2-μ shRNAs (indicated by BFP expression) can reduce the appearance of AP2 complexes in U2OS cells and the accumulation of AP2 complex at the ends of vitronectin-coated nanobars. **c,** Left: Representative fluorescence images showing that knockdown of FCHo1/2 reduces the β5-GFP accumulation at vitronectin-coated nanobar ends. Right: Quantifications of the β5-GFP curvature preference in U2OS cells transfected with scramble or FCHo1/2 shRNAs. N = 12 cells for each condition with each cell represents the average of over 100 nanopillars. Data are mean ± SD. *P* values calculated using t-test. Scale bars: 10 µm.

**Extended Data Fig. 10:**
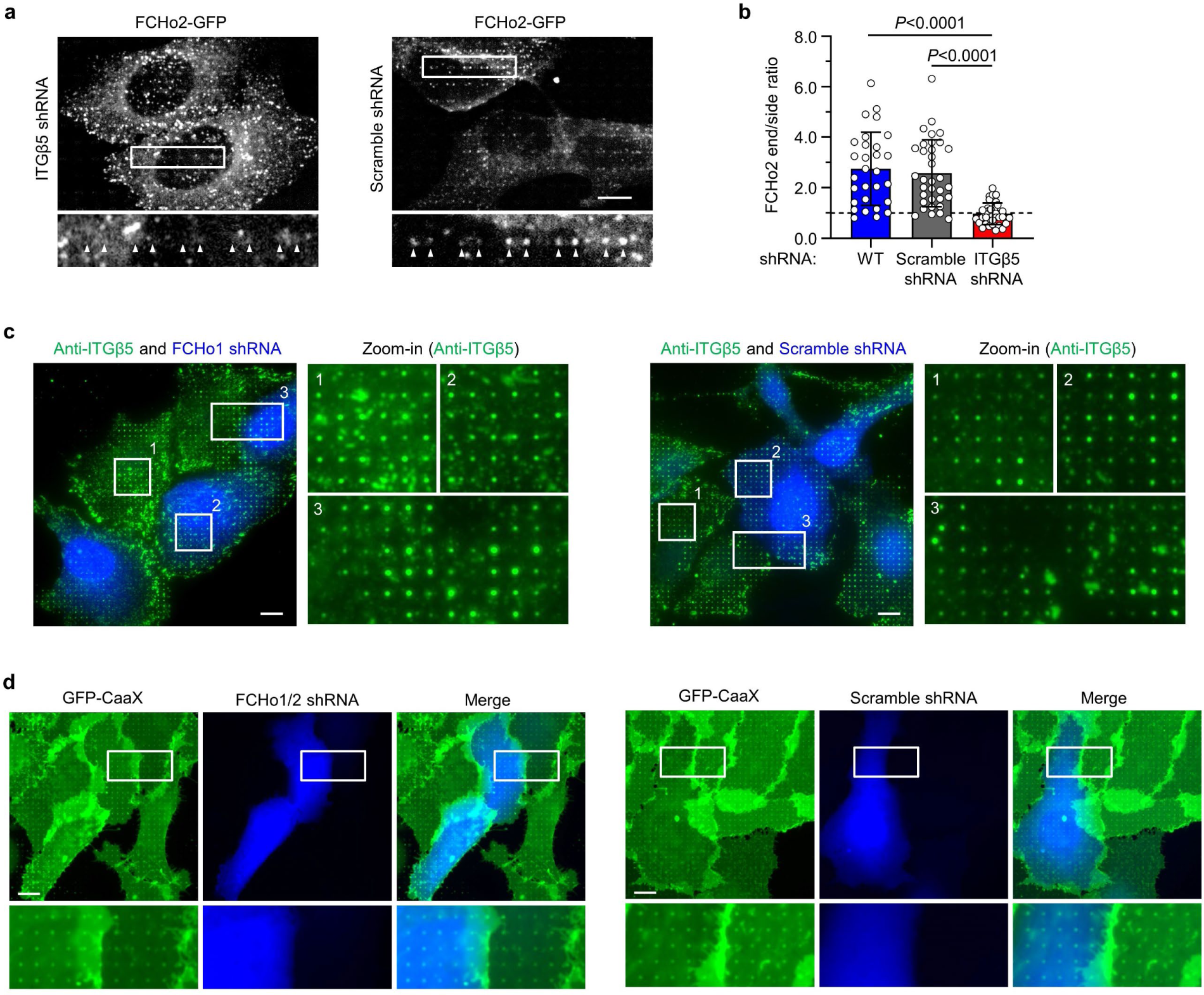
ITGβ5 knockdown reduces the FCHo2 accumulation at vitronectin-coated nanobar ends. **a**, Representative fluorescence images showing that compared with scramble shRNA transfection (right), shRNA knockdown of ITGβ5 (left) reduces the FCHo2-GFP accumulation at the ends of vitronectin-coated nanobars in U2OS cells. **b,** Quantifications of the FCHo2-GFP curvature preference in wild-type (N=29), scramble shRNA-transfected (N=34), or ITGβ5-knockdown (N=36) U2OS cells. Data are mean ± SD. *P* values calculated using one-way ANOVA with Tukey’s multiple comparison. **c,** Representative fluorescence images showing that neither shRNA knockdown of FCHo1 (left) nor scramble shRNA transfection (right) affects ITGβ5 accumulation at vitronectin-coated nanopillars. **d,** Representative fluorescence images showing that compared with scramble shRNA transfection (right), shRNA knockdown of FCHo1/2 (left) does not affect membrane wrapping around vitronectin-coated nanopillars in U2OS cells expressing GFP-CaaX membrane marker. BFP expression is a marker of shRNA transfection. Scale bars: 10 µm.

**Extended Data Fig. 11:**
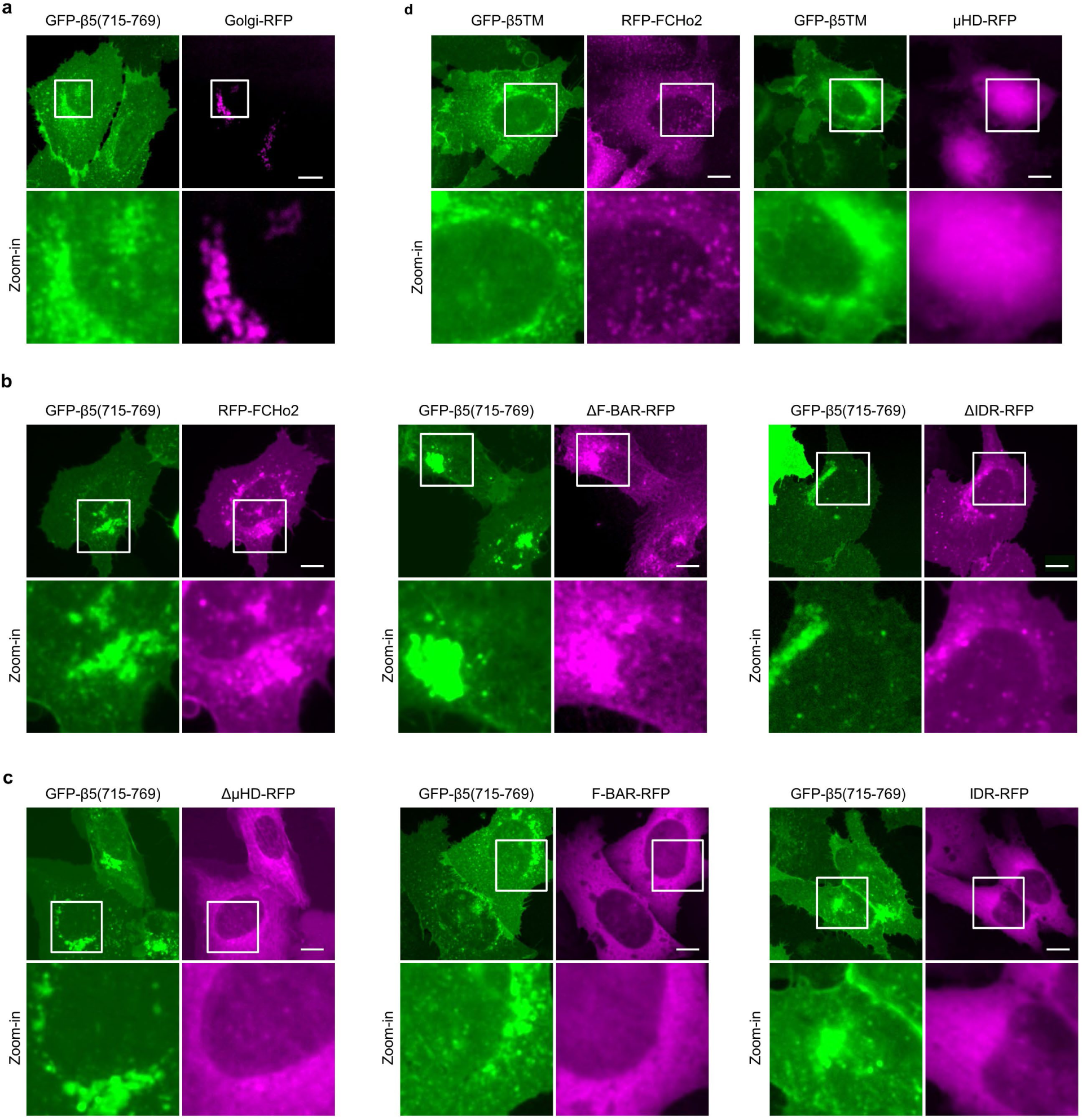
GFP-β5(715-769) induces dramatic redistribution of μHD-containing FCHo2 variants to the plasma membrane and the Golgi apparatus. **a,** GFP-β5(715-769) is located on the plasma membrane with some accumulations around the perinuclear region that colocalizes with Golgi apparatus marker Golgi-RFP. **b,** When co-expressed with GFP-β5(715-769), three μHD domain-containing FCHo2 variants (RFP-FCHo2, FCHo2_ΔF-BAR-RFP, and FCHo2_ΔIDR-RFP) are redistributed to the plasma membrane and colocalize with GFP-β5(715-769) in the perinuclear region. **c,** When co-expressed with GFP-β5(715-769), three FCHo2 variants that don’t contain μHD domain (FCHo2_ΔµHD-RFP, FCHo2_F-BAR-RFP and FCHo2_IDR-RFP) are highly diffusive in the cytosol. **d,** When co-expressed with the negative control GFP-β5TM, RFP-FCHo2 shows cytosolic diffusive pattern with small puncta (left) and FCHo2_µHD is highly cytosolic (right and in Fig. 4l). Scale bars: 10 µm.

**Extended Data Fig. 12:**
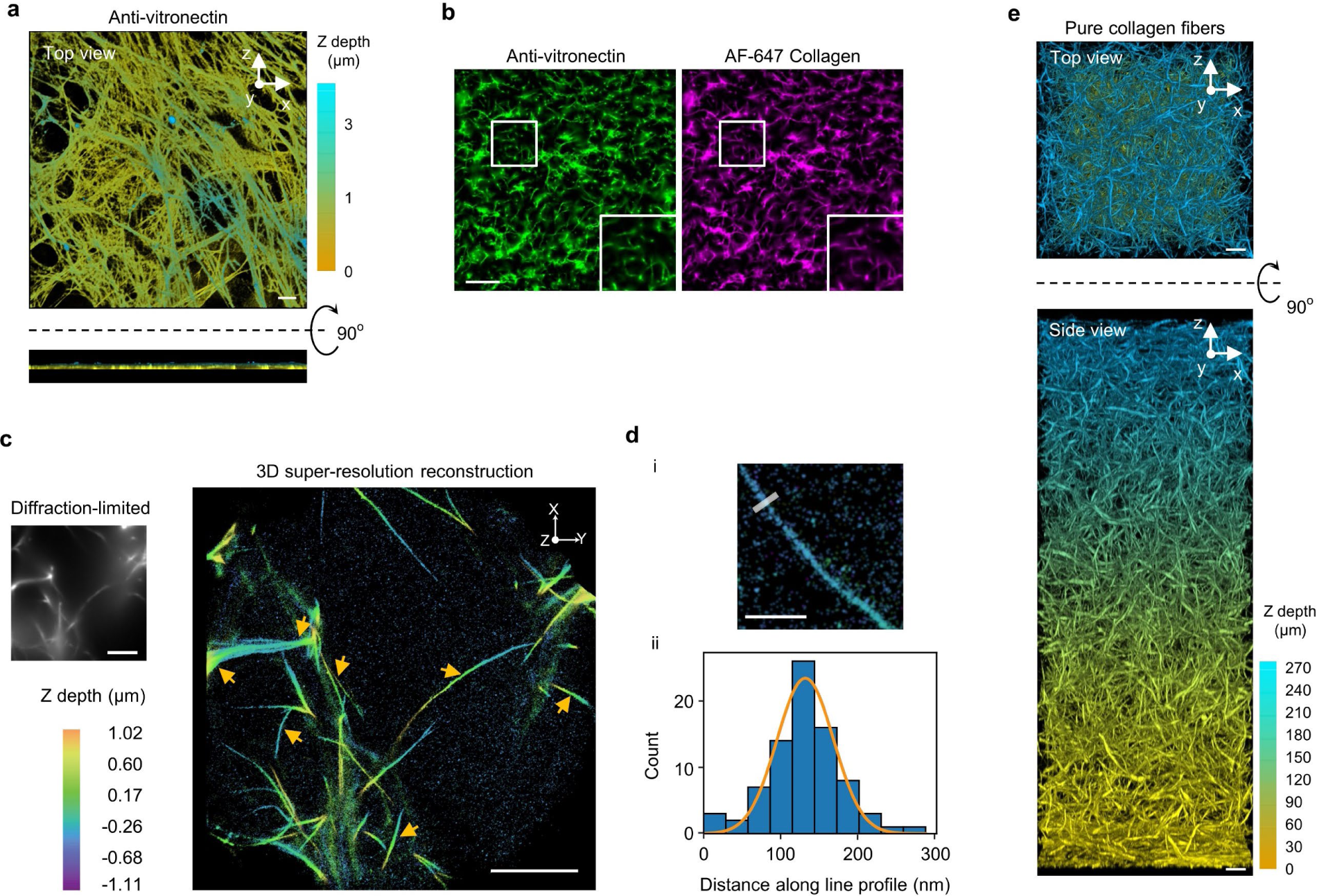
Characterizations of cell-derived ECM, 3D ECM, and individual fibres. **a,** Anti-vitronectin staining illustrates the fibre morphology of IMR-90 lung fibroblast-derived ECM. Z-depth is colour-coded. These cell-derived fibres are dense and flat (< 1 μm thickness) on the glass surface. **b,** In 3D ECM made of vitronectin fibres, colocalization of AF647-collagen with immunolabeled vitronectin confirmed the incorporation of vitronectin in collagen fibres. Scale bar: 50 µm. **c,** 3D super-resolution reconstruction of AF 647-collagen fibres. The widefield diffraction-limited image (top left) depicts collagen fibres. The Z projection of the 3D super-resolution reconstruction (right) reveals higher resolution structures of the collagen fibres in the diffraction-limited image. Yellow arrows point to some individual collagen fibres. Colour encodes the z position. Scale bars: 5 µm. **d,** Extracting the diameter of individual fibres. (i) The image shows an example of a resolved individual fibre. Transverse line profiles (representative example shown at top) are taken across the fibre to estimate its diameter. Scale bar, 1 µm. (ii) Localizations from the line profile are binned into a histogram which is then fit to a Gaussian function (orange line). The full width at half maximum is extracted to estimate the diameter. The line profile in (i) yields the histogram and the fitting is depicted in (ii). **e,** Representative fluorescence images (3D projection, top and side views) of a thick layer (∼ 270 μm) of 3D ECM made of pure collagen fibres labelled with AF647-collagen. Z depth is colour coded. Scale bars: 10 µm.

**Extended Data Fig. 13:**
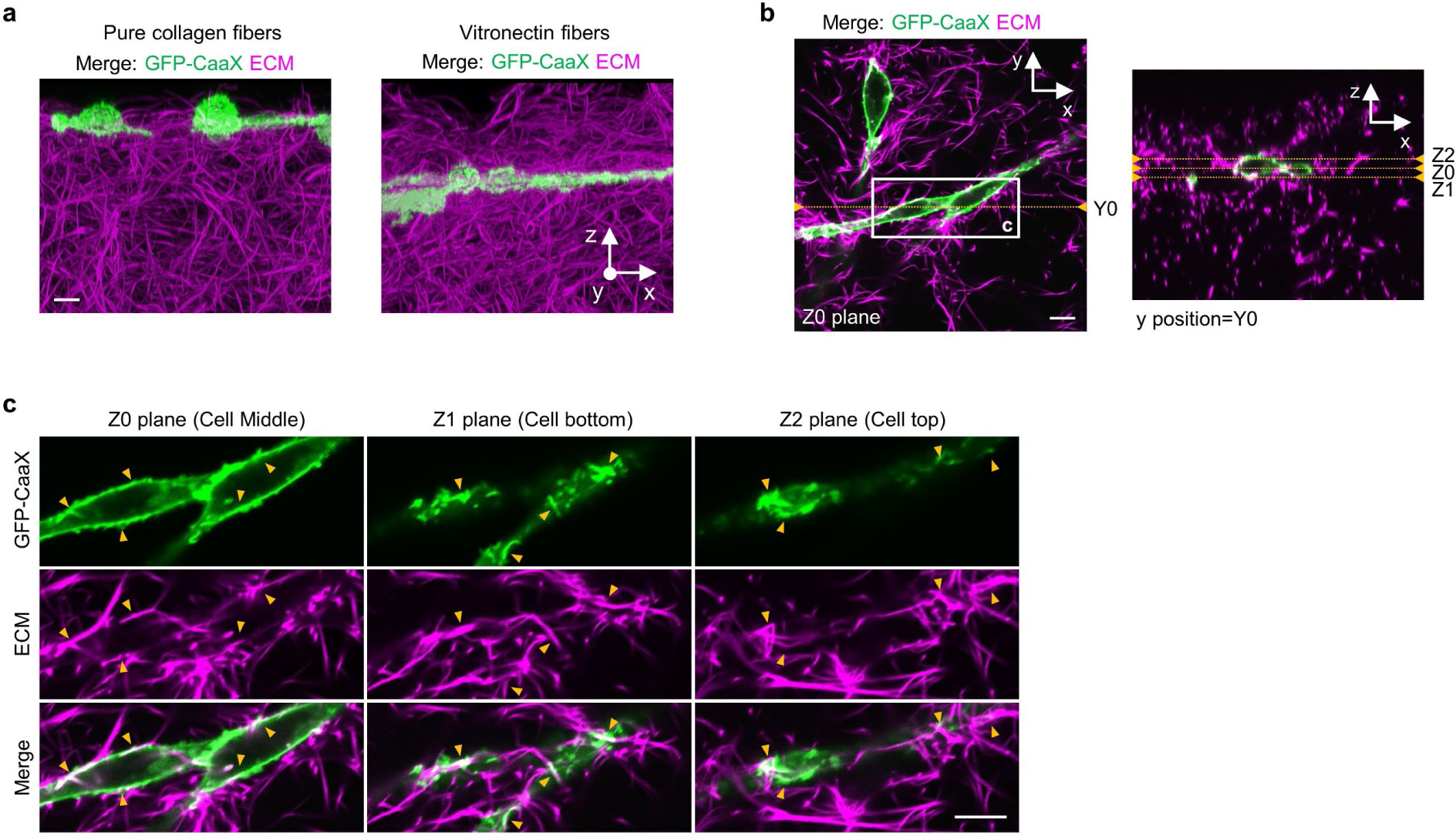
Cell plasma membranes are deformed by 3D ECM fibres. **a,** Representative 3D images (x-z projection) of U2OS cells expressing GFP-CaaX, 72 hrs after being plated on the top of a matrix made of pure collagen fibres (left) or vitronectin fibres (right). **b,** x-y image (Z0 plane) and x-z image (Y0 plane) of Z-stack images of U2OS cells expressing GFP-CaaX in 3D ECM made of vitronectin fibres labelled with AF647-collagen. The 3D projection (side view) has been shown in **a**. **c,** Zoom-in x-y images of the area indicated in **b** showing plasma membrane folding along vitronectin fibres at the middle (z = Z0), bottom (z = Z1), and top (z = Z2) of cells. Scale bars: 10 μm.

**Extended Data Fig. 14:**
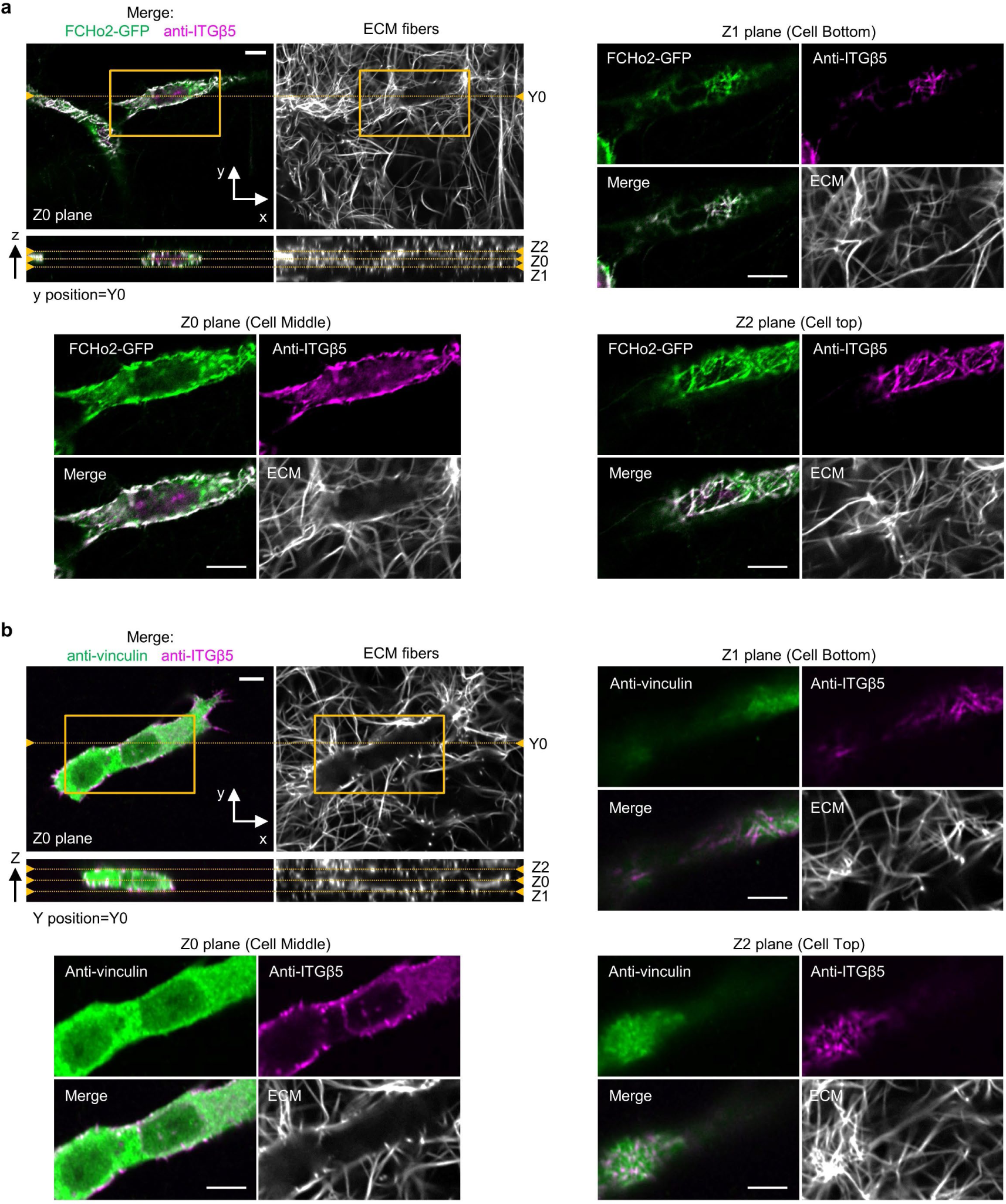
Curved adhesions, but not focal adhesions, are prevalent in 3D ECMs. **a,** x-y view (Z0 plane) and x-z view (Y0 plane) of Z-stack images of immunolabeled ITGβ5 in U2OS cells expressing FCHo2-GFP. The cells are embedded in 3D ECM made of vitronectin fibres labelled with AF647-collagen. Zoom-in x-y images of the yellow-box area show abundant curved adhesions indicated by the colocalization of FCHo2 and ITGβ5. Curved adhesions form along vitronectin fibres at the middle (Z0 plane), bottom (Z1 plane), and top (Z2 plane) of cells. The zoom-in of the x-y slide at the Z0 plane is also shown in Fig. 5g (top). **b,** x-y view (Z0 plane) and x-z view (Y0 plane) of Z-stack images of immunolabeled vinculin and ITGβ5 in U2OS. The cells are embedded in 3D ECM made of vitronectin fibres labelled with AF647-collagen. Zoom-in x-y images of the yellow-box area do not show clear focal adhesions indicated by the colocalization of vinculin and ITGβ5. Examination of different imaging planes, the middle (Z0 plane), bottom (Z1 plane), and top (Z2 plane) of cells show that the colocalization of vinculin and ITGβ5 is sparse. A zoom-in of the x-y view (Z0 plane) is also shown in Fig. 5g (bottom). Scale bars: 10 µm.

**Extended Data Fig. 15:**
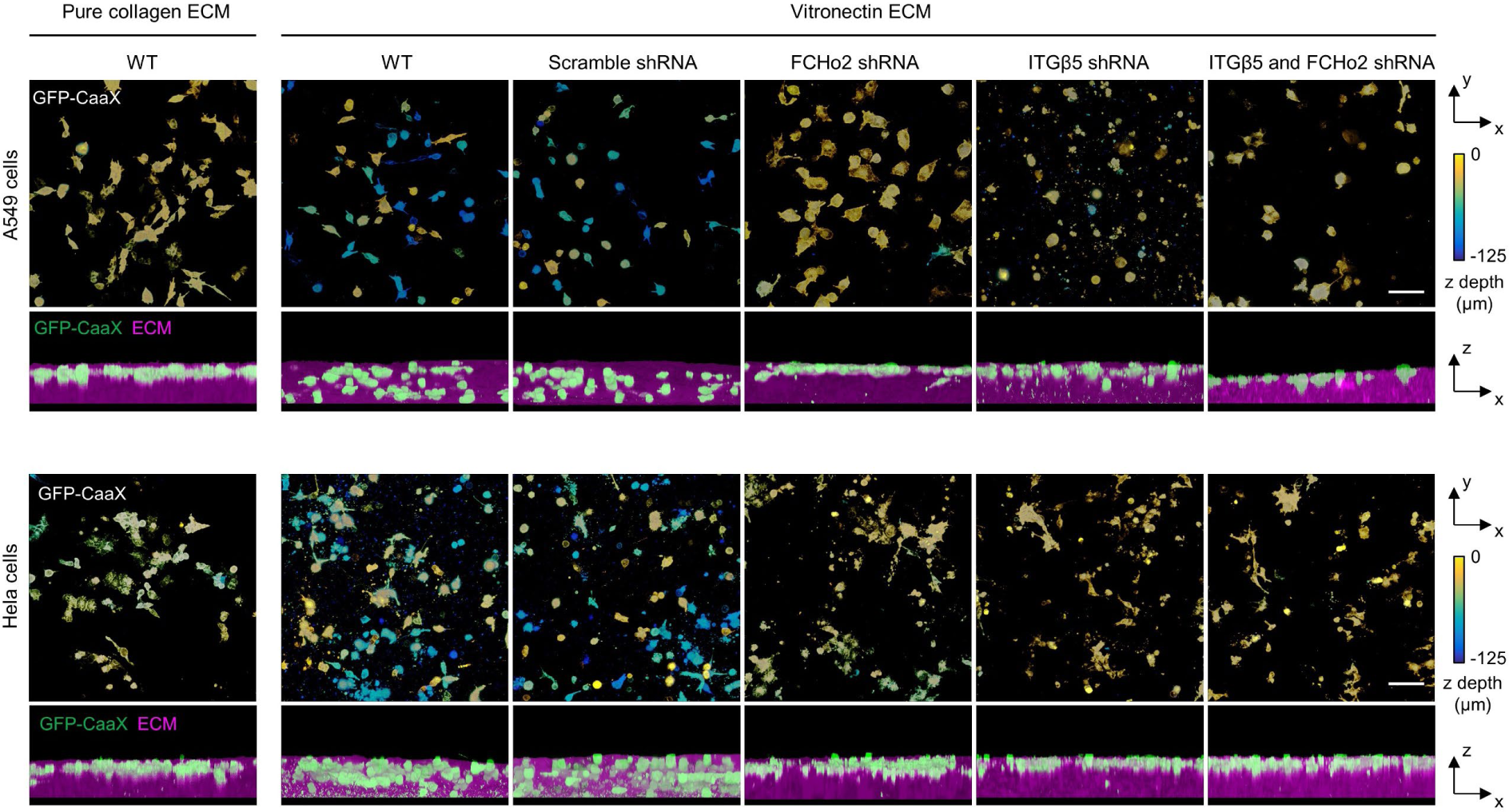
Curved adhesions facilitate cell migration in 3D ECMs. Representative 3D images of A549 cells (top) and HeLa cells (bottom) in 3D matrices after 72-hr culture. Cells were transduced to express GFP-CaaX via lentiviral infection. In the x-y projections, cells are colour-coded according to their depth in the matrix. In the x-z projections, cells are coloured in green and merged with the ECM (magenta). Cells can infiltrate into 3D matrices of vitronectin fibres, but not into 3D matrices of pure collagen fibres. The shRNA knockdown of FCHo2, ITGβ5, or both significantly inhibit cell infiltrations into 3D ECMs. Scale bars: 100 µm.

## Notes

### Competing Interest Statement

The authors have declared no competing interest.

## References

1. Geiger, B., Spatz, J. P. & Bershadsky, A. D. Environmental sensing through focal adhesions. Nat. Rev. Mol. Cell Biol. 10, 21–33 (2009).

2. Kechagia, J. Z., Ivaska, J. & Roca-Cusachs, P. Integrins as biomechanical sensors of the microenvironment. Nat. Rev. Mol. Cell Biol. 20, 457–473 (2019).

3. Harunaga, J. S. & Yamada, K. M. Cell-matrix adhesions in 3D. Matrix Biol. 30, 363–368 (2011).

4. Moser, M., Legate, K. R., Zent, R. & Fässler, R. The tail of integrins, talin, and kindlins. Science 324, 895–899 (2009).

5. Klapholz, B. & Brown, N. H. Talin – the master of integrin adhesions. J. Cell Sci. 130, 2435–2446 (2017).

6. Sun, Z., Costell, M. & Fässler, R. Integrin activation by talin, kindlin and mechanical forces. Nat. Cell Biol. 21, 25–31 (2019).

7. Prager-Khoutorsky, M. et al. Fibroblast polarization is a matrix-rigidity-dependent process controlled by focal adhesion mechanosensing. Nat. Cell Biol. 13, 1457–1465 (2011).

8. Hoffman, B. D., Grashoff, C. & Schwartz, M. A. Dynamic molecular processes mediate cellular mechanotransduction. Nature 475, 316–323 (2011).

9. Case, L. B. & Waterman, C. M. Integration of actin dynamics and cell adhesion by a three-dimensional, mechanosensitive molecular clutch. Nat. Cell Biol. 17, 955–963 (2015).

10. Hakkinen, K. M., Harunaga, J. S., Doyle, A. D. & Yamada, K. M. Direct comparisons of the morphology, migration, cell adhesions, and actin cytoskeleton of fibroblasts in four different three-dimensional extracellular matrices. Tissue Eng. Part A 17, 713–724 (2011).

11. Kubow, K. E. & Horwitz, A. R. Reducing background fluorescence reveals adhesions in 3D matrices. Nature Cell Biology vol. 14 1344–1344 Preprint at https://doi.org/10.1038/ncb2630 (2012).

12. Fraley, S. I. et al. A distinctive role for focal adhesion proteins in three-dimensional cell motility. Nat. Cell Biol. 12, 598–604 (2010).

13. Fraley, S. I., Feng, Y., Wirtz, D. & Longmore, G. D. Reply: reducing background fluorescence reveals adhesions in 3D matrices. Nature Cell Biology vol. 13 5–7 Preprint at https://doi.org/10.1038/ncb0111-5 (2011).

14. Elosegui-Artola, A. et al. Mechanical regulation of a molecular clutch defines force transmission and transduction in response to matrix rigidity. Nat. Cell Biol. 18, 540–548 (2016).

15. Viji Babu, P. K., Rianna, C., Mirastschijski, U. & Radmacher, M. Nano-mechanical mapping of interdependent cell and ECM mechanics by AFM force spectroscopy. Sci. Rep. 9, 12317 (2019).

16. Doyle, A. D. & Yamada, K. M. Mechanosensing via cell-matrix adhesions in 3D microenvironments. Exp. Cell Res. 343, 60–66 (2016).

17. Lock, J. G., et al. Clathrin-containing adhesion complexes. J. Cell Biol. 218, 2086–2095 (2019).

18. Baschieri, F. et al. Frustrated endocytosis controls contractility-independent mechanotransduction at clathrin-coated structures. Nat. Commun. 9, 3825 (2018).

19. Zuidema, A. et al. Mechanisms of integrin αVβ5 clustering in flat clathrin lattices. J. Cell Sci. 131, (2018).

20. Elkhatib, N. et al. Tubular clathrin/AP-2 lattices pinch collagen fibers to support 3D cell migration. Science 356, (2017).

21. Lock, J. G. et al. Reticular adhesions are a distinct class of cell-matrix adhesions that mediate attachment during mitosis. Nat. Cell Biol. 20, 1290–1302 (2018).

22. Vogel, V. & Sheetz, M. Local force and geometry sensing regulate cell functions. Nat. Rev. Mol. Cell Biol. 7, 265–275 (2006).

23. Li, X. et al. A nanostructure platform for live-cell manipulation of membrane curvature. Nat. Protoc. 14, 1772–1802 (2019).

24. Zhao, W. et al. Nanoscale manipulation of membrane curvature for probing endocytosis in live cells. Nat. Nanotechnol. 12, 750–756 (2017).

25. Humphries, J. D., Byron, A. & Humphries, M. J. Integrin ligands at a glance. Journal of Cell Science vol. 119 3901–3903 Preprint at https://doi.org/10.1242/jcs.03098 (2006).

26. Thul, P. J. et al. A subcellular map of the human proteome. Science 356, (2017).

27. Kirchhofer, D., Grzesiak, J. & Pierschbacher, M. D. Calcium as a potential physiological regulator of integrin-mediated cell adhesion. J. Biol. Chem. 266, 4471–4477 (1991).

28. Austen, K. et al. Extracellular rigidity sensing by talin isoform-specific mechanical linkages. Nat. Cell Biol. 17, 1597–1606 (2015).

29. Ringer, P. et al. Multiplexing molecular tension sensors reveals piconewton force gradient across talin-1. Nat. Methods 14, 1090–1096 (2017).

30. Ylänne, J. et al. Distinct functions of integrin alpha and beta subunit cytoplasmic domains in cell spreading and formation of focal adhesions. J. Cell Biol. 122, 223–233 (1993).

31. Yu, C.-H. et al. Integrin-beta3 clusters recruit clathrin-mediated endocytic machinery in the absence of traction force. Nat. Commun. 6, 8672 (2015).

32. Nakamoto, M. L., Forró, C., Zhang, W., Tsai, C.-T. & Cui, B. Expansion Microscopy for Imaging the Cell–Material Interface. ACS Nano 16, 7559–7571 (2022).

33. Henne, W. M. et al. FCHo proteins are nucleators of clathrin-mediated endocytosis. Science 328, 1281–1284 (2010).

34. Horton, E. R. et al. Definition of a consensus integrin adhesome and its dynamics during adhesion complex assembly and disassembly. Nat. Cell Biol. 17, 1577–1587 (2015).

35. Hollopeter, G., et al. The membrane-associated proteins FCHo and SGIP are allosteric activators of the AP2 clathrin adaptor complex. eLife vol. 3 Preprint at https://doi.org/10.7554/elife.03648 (2014).

36. Berman, J., Stoner, G., Dawe, C., Rice, J. & Kingsbury, E. Histochemical demonstration of collagen fibers in ascorbic-acid-fed cell cultures. In Vitro 14, 675–685 (1978).

37. Hayman, E. G., Pierschbacher, M. D., Ohgren, Y. & Ruoslahti, E. Serum spreading factor (vitronectin) is present at the cell surface and in tissues. Proc. Natl. Acad. Sci. U. S. A. 80, 4003– 4007 (1983).

38. Gebb, C., Hayman, E. G., Engvall, E. & Ruoslahti, E. Interaction of vitronectin with collagen. Journal of Biological Chemistry vol. 261 16698–16703 Preprint at https://doi.org/10.1016/s0021-9258(18)66621-9 (1986).

39. Doyle, A. D. Fluorescent labeling of rat-tail collagen for 3D fluorescence imaging. Bio Protoc. 8, (2018).

40. Ishikawa, M. & Hayashi, M. Activation of the collage-binding of endogenous serum vitronectin by heating, urea and glycosaminoglycans. Biochimica et Biophysica Acta (BBA) - Protein Structure and Molecular Enzymology 1121, 173–177 (1992).

41. Zhang, L.-Y. et al. Integrin Beta 5 Is a Prognostic Biomarker and Potential Therapeutic Target in Glioblastoma. Front. Oncol. 9, 904 (2019).

42. Bianchi-Smiraglia, A., Paesante, S. & Bakin, A. V. Integrin β5 contributes to the tumorigenic potential of breast cancer cells through the Src-FAK and MEK-ERK signaling pathways. Oncogene vol. 32 3049–3058 Preprint at https://doi.org/10.1038/onc.2012.320 (2013).

43. Hurtado de Mendoza, T., et al. Tumor-penetrating therapy for β5 integrin-rich pancreas cancer. Nat. Commun. 12, 1541 (2021).

44. Boyd, N. A., Bradwell, A. R. & Thompson, R. A. Quantitation of vitronectin in serum: evaluation of its usefulness in routine clinical practice. J. Clin. Pathol. 46, 1042–1045 (1993).

45. Stathakis, N. E., Fountas, A. & Tsianos, E. Plasma fibronectin in normal subjects and in various disease states. J. Clin. Pathol. 34, 504–508 (1981).

46. Gospodarowicz, D., Gonzalez, R. & Fujii, D. K. Are factors originating from serum, plasma, or cultured cells involved in the growth-promoting effect of the extracellular matrix produced by cultured bovine corneal endothelial cells? J. Cell. Physiol. 114, 191–202 (1983).

47. Pavani, S. R. P. et al. Three-dimensional, single-molecule fluorescence imaging beyond the diffraction limit by using a double-helix point spread function. Proc. Natl. Acad. Sci. U. S. A. 106, 2995–2999 (2009).

48. Roy, A. R. et al. Exploring cell surface-nanopillar interactions with 3D super-resolution microscopy. ACS Nano (2021) doi:10.1021/acsnano.1c05313.

49. Lu, C.-H. et al. Membrane curvature regulates the spatial distribution of bulky glycoproteins. Nat. Commun. 13, 3093 (2022).

50. Lou, H.-Y. et al. Membrane curvature underlies actin reorganization in response to nanoscale surface topography. Proc. Natl. Acad. Sci. U. S. A. 116, 23143–23151 (2019).

