## Supplementary Information for "Curved adhesions mediate cell attachment to soft matrix fibres in 3D"

### Supplementary Table 1: Information on shRNAs.

| Target protein | Source | Identifier | Target sequence | KD efficiency |
|---|---|---|---|---|
| Integrin β5 | Sigma-Aldrich | TRCN0000296117<br>TRCN0000289121<br>TRCN0000296116 | AGCTTGTTGTCCCAATGAAAT<br>CTGAGGGCAAACCTTGTCAAA<br>GGATCAGCCTGAGGATCTTAA | 85%<br>84%<br>82% |
| AP2-μ | Sigma-Aldrich | TRCN0000060239<br>TRCN0000060241 | CACCAGCTTCTTCCACGTTAA<br>GCTGGATGAGATTCTAGACTT | 94%<br>95% |
| Clathrin heavy chain | Sigma-Aldrich | TRCN0000011216<br>TRCN0000007981 | CGGTTGCTCTTGTTACGGATA<br>CGTGTTCTTGTAACCTTTATT | 88%<br>97% |
| FCHo1 | Sigma-Aldrich | TRCN0000162794<br>TRCN0000164648 | CACAACCGCTATTGAGCACTT<br>GCAGGAAGCGATGAAACGTTT | NA<br>NA |
| FCHo2 | Sigma-Aldrich | TRCN0000167218<br>TRCN0000167925 | GCTACAGTATTAAACCAGAAA<br>CCAAAGCTTACTTCAGGCAAA | 88%<br>75% |
| EPS15 | Sigma-Aldrich | TRCN0000007980<br>TRCN0000007978 | CCCAGAATGGATTGGAAGTTT<br>GCAGTGAAACAGCCAACCTTA | 86%<br>79% |
| EPS15R | Sigma-Aldrich | TRCN0000233084<br>TRCN0000233083 | GAGCATGCCACCGCCTAAATT<br>AGTCTGGCCTCTCGGACATTA | 77%<br>72% |
| Intersectin 1 | Sigma-Aldrich | TRCN0000002009<br>TRCN0000002010 | GCACTAGCTGACATGAATAAT<br>GCAGTTGTTTGATGAGCCGTA | 64%<br>56% |
| Intersectin 2 | Sigma-Aldrich | TRCN0000002385<br>TRCN0000318540 | CCTGGACTGCAAAGAAAGATA<br>GCAGAACGTAAAGCCCAGAAA | 75%<br>84% |
| Scramble (Negative control) | Addgene | Cat#1864 | CCTAAGGTTAAGTCGCCCTCG | NA |

### Supplementary Table 2. Information on antibodies.

| Name | Source | Identifier |
|---|---|---|
| **Primary antibodies** | | |
| Rat anti-integrin β1 antibody (clone 9EG7) | BD Biosciences | Cat#550531 |
| Mouse anti-activated integrin β1 antibody (clone HUTS-4) | Sigma-Aldrich | Cat#MAB2079Z |
| Mouse anti-integrin αvβ5 antibody (clone P5H9) | R&D Systems | Cat#MAB2528 |
| Mouse anti-alpha adaptin antibody [AP6] | Abcam | Cat#ab2730 |
| Mouse anti-vinculin antibody (clone hVIN-1) | Sigma-Aldrich | Cat#V9131 |
| Mouse anti-paxillin antibody (clone 349/Paxillin) | BD Biosciences | Cat#610051 |
| Mouse anti-talin1 antibody (clone 8D4) | Abcam | Cat#ab157808 |
| Mouse Anti-Vitronectin/S-Protein antibody (clone VN58-1) | Abcam | Cat#ab13413 |
| Rabbit anti-FAK (phospho Y397) antibody (clone EP2160Y) | Abcam | Cat#ab81298 |
| Rabbit anti-Clathrin heavy chain antibody (Polyclonal) | Abcam | Cat#ab21679 |
| Rabbit anti-integrin β5 antibody (clone D24A5) | Cell signaling technology | Cat#3629S |
| Rabbit anti-FCHo2 antibody (Polyclonal) | Novus Biologicals | Cat#NBP2-32694 |
| Rabbit anti-GAPDH (clone 14C10) | Cell signaling technology | Cat#2118 |
| Rabbit anti-GFP antibody (Polyclonal) | Invitrogen | Cat#A-11122 |
| Mouse anti-mCherry antibody (clone GT857) | Sigma-Aldrich | Cat#SAB2702291 |
| **Secondary antibodies** | | |
| Alexa Fluor 488-conjugated goat anti-Mouse IgG (H+L) antibody | Thermo Fisher Scientific | Cat#A-11001 |
| Alexa Fluor 568-conjugated goat anti-Mouse IgG (H+L) antibody | Thermo Fisher Scientific | Cat#A-11004 |
| Alexa Fluor 488-conjugated goat anti-Rabbit IgG (H+L) antibody | Thermo Fisher Scientific | Cat#A-11034 |
| Texas Red-conjugated goat anti-Rabbit IgG (H+L) antibody | Thermo Fisher Scientific | Cat#T2767 |
| Alexa Fluor 647-conjugated goat anti-Rat IgG (H+L) antibody | Thermo Fisher Scientific | Cat#A-21247 |
| HRP-linked goat anti-Rabbit IgG (H+L) antibody | Cell signaling technology | Cat#7074 |
| HRP-linked goat anti-Mouse IgG (H+L) antibody | Cell signaling technology | Cat#7076 |

### Supplementary Table S3: The statistics for null hypothesis testing

| Figure | Test | | Degrees of freedom | Specifics |
|---|---|---|---|---|
| Fig. 1k | One-way ANOVA | | F (10,77) = 140.8 | $R^2 = 0.9481$ |
| Fig. 2d | Kruskal-Wallis test | | 3 | H = 1222 |
| Fig. 2i | One-way ANOVA | | F (7,72) = 97.35 | $R^2 = 0.9044$ |
| Fig. 2m | Kruskal-Wallis test | Vitronectin | 5 | H = 91.28 |
| | | Fibronectin | 5 | H = 67.26 |
| | Mann Whitney test | Gelatin | n/a | U = 720.5 |
| | t-test | PLL | t = 0.1411, df = 70 | $R^2 = 0.0002843$ |
| | t-test | BSA | t = 1.246, df = 73 | $R^2 = 0.02083$ |
| Fig. 3d | One-way ANOVA | | F (4, 55) = 0.4478 | $R^2 = 0.03154$ |
| Fig. 3i | t-test | | t = 7.761, df = 17.07 | $R^2 = 0.7792$ |
| | One-way ANOVA | | F (4, 55) = 120.3 | $R^2 = 0.8974$ |
| Fig. 4d | Mann Whitney test | | n/a | U = 2998 |
| Fig. 4g | One-way ANOVA | | F (3, 181) = 39.93 | $R^2 = 0.3983$ |
| Fig. 4j | One-way ANOVA | | F (8, 225) = 60.44 | $R^2 = 0.6824$ |
| Fig. 5h | Kruskal-Wallis test | Number | 3 | H = 45.30 |
| | t-test | Size 2D | t = 13.61, df = 21 | $R^2 = 0.8982$ |
| | | Size 3D | t = 5.423, df = 8 | $R^2 = 0.7861$ |
| Fig. 6c | Kruskal-Wallis test | U2OS | 5 | H = 294.2 |
| | | A549 | 5 | H = 203.6 |
| | | Hela | 5 | H = 113.3 |
| Extended Data Fig. 3c | Kruskal-Wallis test | | 3 | H = 969.4 |
| Extended Data Fig. 7e | Kruskal-Wallis test | | 2 | H = 58.3 |
| Extended Data Fig. 9c | t-test | | t = 11.76, df = 16 | $R^2 = 0.8963$ |
| Extended Data Fig. 10b | One-way ANOVA | | F (2, 96) = 25.46 | $R^2 = 0.3466$ |
| *Confidence level is 0.05 for all tests. | | | | |

**Supplementary Video 1. Dynamics of curved adhesions at nanopillars.**

Live-cell fluorescence imaging of U2OS cell expressing ITGβ5-GFP stained with a plasma membrane marker CellMask Red on vitronectin-coated nanopillars at 15 s/frame for ~80 min. The ratiometric images of ITGβ5/membrane (normalized by its mean per cell) are shown in the Parula color scale. Time (in seconds) is indicated. Most curved adhesions at nanopillars are stable for the ~80-min time duration. Meanwhile, few curved adhesions slowly assemble and disassemble at nanopillars. Scale bar, 5 μm.

**Supplementary Video 2. Dynamics of FCHo2 accumulations in curved adhesions at nanopillars.**

Live-cell fluorescence imaging of U2OS cell co-expressing ITGβ5-GFP and RFP-FCHo2 on vitronectin-coated nanopillars at 15 s/frame for ~20 min. Time (in seconds) is indicated. The FCHo2 accumulations are strong and stable in curved adhesions marked by ITGβ5 accumulations at nanopillars (representative ones are indicated by white circles), suggesting that FCHo2 is an integral component of curved adhesions. Some nanopillars without ITGβ5 accumulation (representative ones indicated by yellow circles) also show FCHo2 accumulations, but these accumulations are usually weak and dynamic, exhibiting frequent assembly and disassembly on a time scale of minutes. Scale bar, 5 μm.

**Supplementary Video 3. Z-stack images of cells staying on the top of a 3D matrix of pure collagen fibers.**

Confocal fluorescence images of U2OS cells expressing GFP-CaaX (green). Cells were plated on the top of a 3D matrix made of collagen fibers labeled with AF647-collagen. After 72 hours of culture, cells mostly stay on the top of the matrix (magenta). The top 100 μm of the sample was imaged every 0.5 μm from the bottom to the top. The Z positions are indicated. Scale bar, 10 μm. The corresponding 3D projection (side view) is shown in **Extended Data Fig. 13a** (left).

**Supplementary Video 4. Z-stack images of cells embedded in a 3D matrix of vitronectin fibers.**

Confocal fluorescence images of U2OS cells expressing GFP-CaaX (green). Cells were plated on the top of a 3D matrix made of vitronectin fibers labeled with AF647-collagen. After 72 hours of culture, cells have infiltrated and are fully embedded in the matrix (magenta). The top 100 μm of the sample was imaged every 0.5 μm from the bottom to the top. The Z positions are indicated. Scale bar, 10 μm. The corresponding 3D projection (side view) is shown in **Extended Data Fig. 13a** (right).

**Supplementary Video 5. Z-stack images of curved adhesions in 3D.**

Confocal fluorescence images of immunolabeled ITGβ5 (magenta) in U2OS cells expressing FCHo2-GFP (green) embedded in a 3D matrix made of vitronectin fibers labeled with AF647-collagen (grayscale). Curved adhesions, the colocalizations of ITGβ5 and FCHo2 along vitronectin fibers, form extensively. The images were acquired every 0.5 μm from the bottom to the top and the Z positions are indicated. Scale bar, 25 μm.

**Supplementary Video 6. Z-stack images of focal adhesions in 3D.**

Confocal fluorescence images of immunolabeled vinculin (green) and ITGβ5 (magenta) in U2OS cells embedded in a 3D matrix made of vitronectin fibers labeled with AF647-collagen (grayscale). Focal adhesions, the colocalizations of ITGβ5 and vinculin, are barely observed throughout the cells. Meanwhile, ITGβ5 still extensively accumulates along the ECM fibers. The images were acquired every 0.5 μm from the bottom to the top and the Z positions are indicated. Scale bar, 25 μm.
